# Dynamics of RNA localization to nuclear speckles are connected to splicing efficiency

**DOI:** 10.1101/2024.02.29.581881

**Authors:** Jinjun Wu, Yu Xiao, Yunzheng Liu, Li Wen, Chuanyang Jin, Shun Liu, Sneha Paul, Chuan He, Oded Regev, Jingyi Fei

## Abstract

Nuclear speckles, a type of membraneless nuclear organelle in higher eukaryotic cells, play a vital role in gene expression regulation. Using the reverse transcription-based RNA-binding protein binding sites sequencing (ARTR-seq) method, we study human transcripts associated with nuclear speckles. We identify three gene groups whose transcripts demonstrate different speckle localization properties and dynamics: stably enriched in nuclear speckles, transiently enriched in speckles at the pre-mRNA stage, and not enriched in speckles. Specifically, we find that stably-enriched transcripts contain inefficiently spliced introns. We show that nuclear speckles specifically facilitate splicing of speckle-enriched transcripts. We further reveal RNA sequence features contributing to transcript speckle localization, underscoring a tight interplay between genome organization, RNA cis-elements, and transcript speckle enrichment, and connecting transcript speckle localization with splicing efficiency. Finally, we show that speckles can act as hubs for the regulated retention of introns during cellular stress. Collectively, our data highlight a role of nuclear speckles in both co– and post-transcriptional splicing regulation.

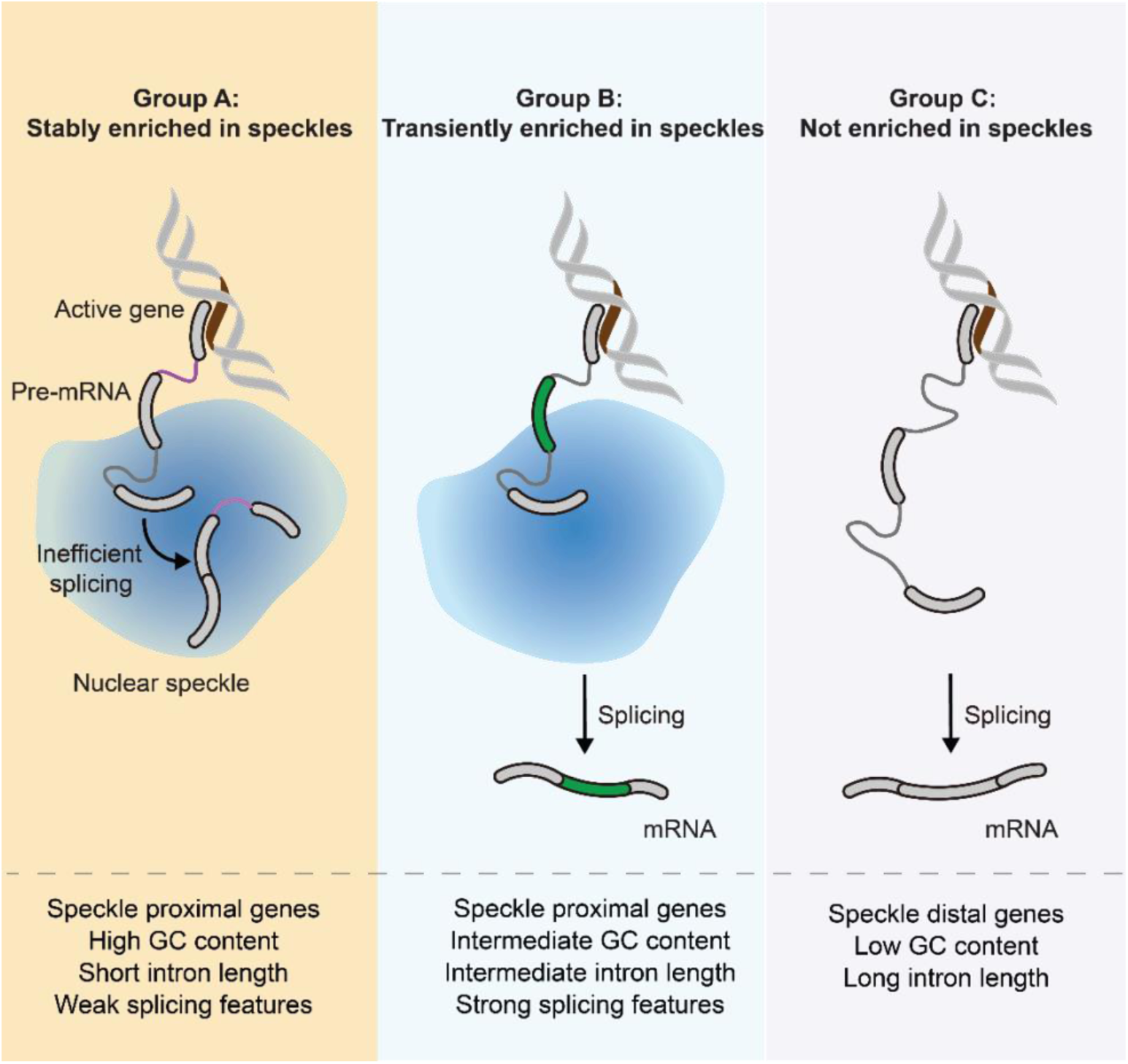
GRAPHICAL ABSTRACT.

## INTRODUCTION

Membraneless organelles are prevalent in eukaryotic cells and are broadly involved in processing and assembly of ribonucleoprotein complexes, gene expression, signal transduction, and stress responses (Banani et al., 2017; Hirose et al., 2023). Nuclear speckles are a type of membraneless organelles in the nucleus of higher eukaryotic cells. A typical cell contains tens of nuclear speckles, ranging in size from a few hundred nanometers to a few microns. Nuclear speckles have a rich proteome consisting of over a hundred protein species, many of which are spliceosomal components or splicing regulators (Galganski et al., 2017; Ilık and Aktaş, 2022). Their core region is defined by the two scaffold proteins SON and SRRM2 (Ilik et al., 2020). Nuclear speckles also have a rich transcriptome, including polyadenylated (polyA) RNAs, and certain long noncoding RNAs (lncRNAs), as was recently systematically mapped through APEX-seq (Barutcu et al., 2022). Changes in nuclear speckle composition or morphology are frequently associated with cancers, neuronal disorders and infection (Galganski et al., 2017; Lester et al., 2021; Mor et al., 2016; Sellier et al., 2010; Urbanek et al., 2016). But the fundamental roles of nuclear speckles in gene expression remain elusive, making it challenging to mechanistically connect nuclear speckles to pathogenesis of these diseases.

While nuclear speckles were historically considered as storage sites for splicing factors, new evidence now portrays nuclear speckles as active hubs promoting gene expression (Galganski et al., 2017; Ilık and Aktaş, 2022; Chen and Belmont, 2019). A positive correlation has been observed between expression level and the speckle proximity of gene foci, both in fluorescence imaging using in situ hybridization (Ding and Elowitz, 2019; Hall et al., 2006) and in more recent genome-wide sequencing through proximity labeling (W. Chen et al., 2018; Y. Chen et al., 2018; Quinodoz et al., 2018). It was suggested that around 50% of transcriptionally active genes are associated with nuclear speckles (Y. Chen et al., 2018). However, it is unclear why certain genes are associated with speckles while others do not. Gene foci have also been observed to localize close to nuclear speckles in a regulated fashion. For example, α-globin and β-globin genes are localized to nuclear speckles when actively transcribed during erythroid differentiation (Brown et al., 2006), and *HSPA1A* transgenes were observed to move towards nuclear speckles upon heat shock (Khanna et al., 2014). Moreover, as demonstrated with p53, transcription factors can drive localization of a subset of their target genes to nuclear speckles, enhancing their RNA expression levels (Alexander et al., 2021).

Being a compartment enriched in splicing factors (Galganski et al., 2017; Ilık and Aktaş, 2022), nuclear speckles are tightly linked to splicing. Detection within nuclear speckles of phosphorylated SF3b, considered as a marker for active spliceosomes (Bessonov et al., 2010; Hluchý et al., 2022; Shi et al., 2006; Wang et al., 1998), suggests active splicing taking place in speckles (Girard et al., 2012). The enhanced enrichment of polyA RNAs in speckles upon splicing inhibition is indicative of speckles as a compartment to accommodate post-transcriptionally accumulated transcripts (Girard et al., 2012; Kaida et al., 2007). In line with this observation, the recent APEX-seq mapping of nuclear speckle-localized transcriptome revealed an enrichment of retained introns in nuclear speckles (Barutcu et al., 2022). These results suggest that nuclear speckles serve as a post-transcriptional quality control compartment for incompletely spliced transcripts. In addition, nuclear speckles were demonstrated to promote co-transcriptional splicing through increased binding of spliceosomes to pre-mRNAs from speckle-proximal genes (Bhat et al., 2023; Ding and Elowitz, 2019). In addition to promoting constitutive splicing, nuclear speckles were also demonstrated to facilitate splicing of stress-related genes under ribotoxic stress in a regulated fashion (Sung et al., 2023), as well as impacting alternative splicing (Xu et al., 2022). However, within these proposed functions of nuclear speckle in splicing, fundamental questions are not addressed: (1) Do all genes require speckles for co-or post-transcriptional splicing? (2) If not, do transcripts utilizing speckles for co– or post-transcriptional splicing differ in any way? Addressing these questions will provide us a clearer framework of how nuclear speckles coordinate co– and post-transcriptional splicing.

In this work, we employ ARTR-seq (reverse transcription-based RNA-binding protein binding sites sequencing) (Xiao et al., 2024) to comprehensively quantify the speckle transcriptome. We identify three gene groups whose transcripts demonstrate different speckle localization properties: transcripts from Group A genes are stably enriched in nuclear speckles; transcripts from Group B genes are transiently enriched in speckles at the pre-mRNA stage; and transcripts from Group C genes are not enriched in speckles. Through a biochemical assay, we demonstrate a functional role of nuclear speckles in promoting splicing of speckle-associated transcripts from Group A and B genes. We further reveal RNA sequence cis-elements that contribute to transcript speckle localization, suggesting a tight interplay between gene position, sequence features, and transcript speckle enrichment. Finally, using heat shock as an example, we suggest that nuclear speckles act as hubs for the regulated retention of introns during cellular stress.

## RESULTS

### Mapping the nuclear speckle transcriptome

To map the nuclear speckle-enriched transcriptome, we adopted our recently developed method ARTR-seq (Xiao et al., 2024). This method uses a recombinant enzyme consisting of Protein A/G fused to a reverse transcriptase (pAG-RTase) (Figure 1A). Protein A/G combines the IgG binding domains of Protein A and protein G and can thus bind to most IgG subclasses including polyclonal or monoclonal IgG antibodies. Because the physical distance between the RTase and the targeted protein is estimated to be at most ∼45 nm, considering the physical dimension of antibodies (∼14 nm), pAG (∼3 nm), RTase (∼4 nm), and the 30 amino acid linker in between (∼10 nm) (Zou et al., 2019), we reasoned that the method is well suited for identification of the transcriptome within membraneless organelles. Following cell fixation and permeabilization, a nuclear speckle scaffold protein (either SON or SRRM2) was first labeled with primary and secondary antibodies, and then pAG-RTase. Reverse transcription was initiated by exogenous addition of biotin dNTPs and other reaction components, followed by cell lysis, RNase treatment, and affinity enrichment of biotinylated cDNAs using streptavidin beads. The biotinylated cDNAs were ligated with an adaptor and prepared for library amplification and sequencing. The fluorophore-labeled pAG-RTase showed good colocalization with antibodies against SON and biotin-labeled cDNA (Figure 1B), with all signals exhibiting specific nuclear speckle localization. The colocalization analysis confirms that pAG-RTase can effectively perform reverse transcription in situ with the desired spatial localization. Here we denote the number of reads mapping to each gene (whether in exons or in introns) as N_SON_ when SON is targeted and as N_SRRM2_ when SRRM2 is targeted.

**Figure 1.**
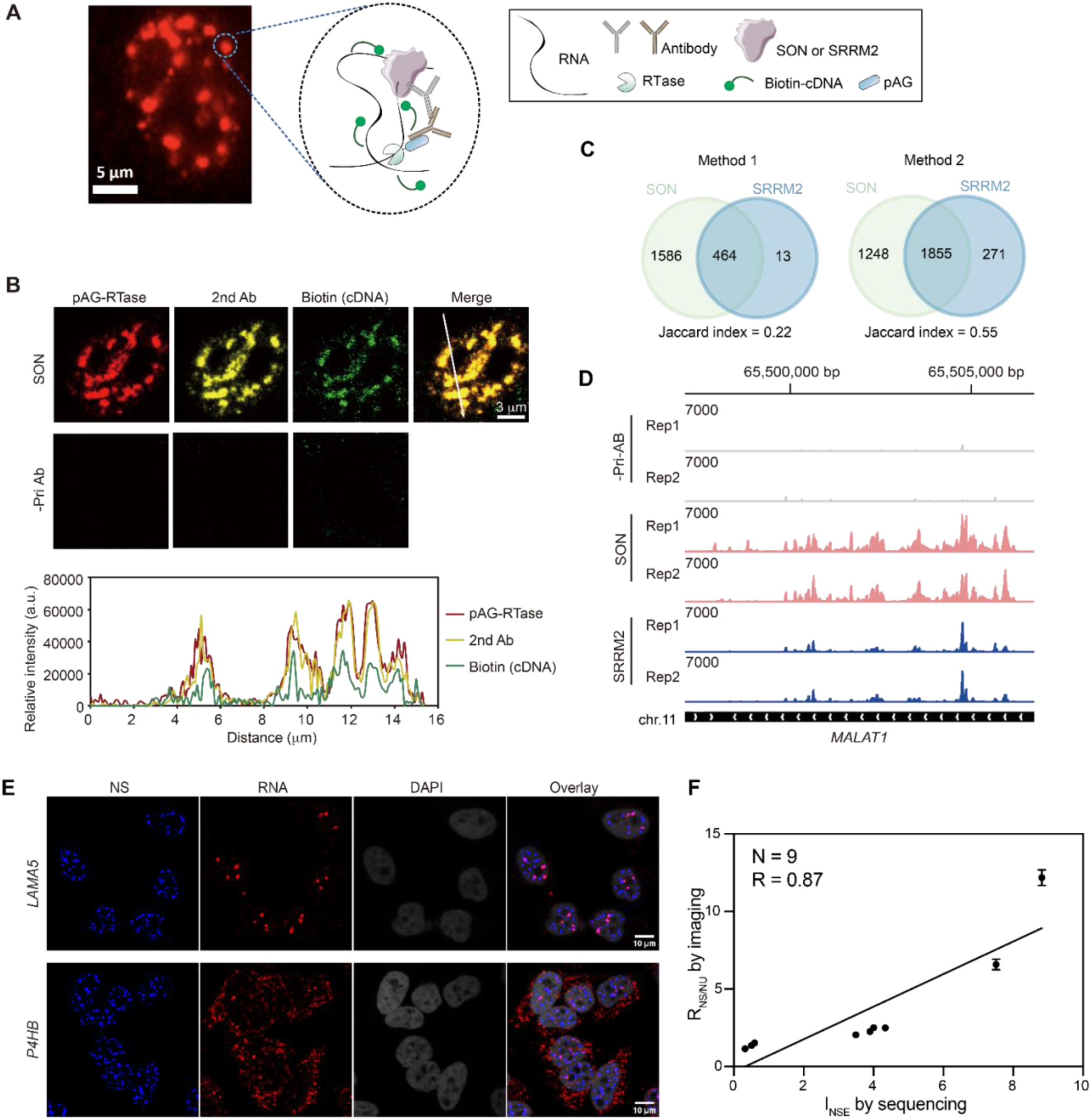
Characterization of nuclear speckle-enriched transcriptome using ARTR-seq. (A) Scheme of ARTR-seq. Specific speckle scaffold protein is immunostained by primary and secondary antibodies sequentially. pAG-RTase is then allowed to bind to the antibody to initiate reverse transcription in situ. The generated biotinylated cDNAs are collected and prepared for sequencing. (B) Representative image showing colocalization of pAG-RTase labeled with Alexa Fluor 647 (AF647) (red), secondary antibody labeled with Alexa Fluor 568 (AF568, yellow) against anti-SON primary antibody, and generated biotinylated-cDNA detected by Alexa Fluor 488 (AF488) labeled antibody against biotin (green). Scale bar: 3 µm. (C) Venn diagram of overlapped speckle-enriched genes identified through targeting SON and SRRM2 in HeLa cells using Method 1 or Method 2. Speckle-enriched transcripts are defined by I_NSE_>2 and adjusted p-value < 0.05. (D) Genome tracks showing ARTR-seq reads generated from targeting SON or SRRM2 proteins, and from control samples without primary antibody, mapped to gene locus encoding lncRNA *MALAT1*. (E) RNA FISH images showing speckle-enriched *LAMA5* transcript in comparison with speckle non-enriched *P4HB* transcript. RNA FISH probes are labeled with AF647 (red). Nucleus was stained with DAPI. Scale bar: 10 μm. (F) Correlation between speckle partition coefficient (R_NS/NU_) measured by RNA FISH imaging and I_NSE_ values determined by ARTR-seq using Method 1.

We included two additional RNA-seq datasets as controls for the calculation of speckle enrichment (Figure S1). The first dataset consists of ARTR-seq obtained from a sample prepared without the primary antibody (with reads denoted by N_-priAB_). Without the primary antibody, the secondary antibody and pAG-RTase exhibited minimal nonspecific binding (Figure 1B). The second dataset comprises rRNA-depleted nuclear RNA-seq (reads denoted by N_nu-RNA_), reflecting the nuclear abundance of each RNA species. Importantly, we observed a strong correlation between N_-priAB_ and N_nu-RNA_ (Figure S2A), supporting that N_-priAB_ can also quantitatively reflect the nuclear RNA abundance. We then calculated an Index for nuclear speckle enrichment (I_NSE_) in two ways. In Method 1, I_NSE_ was calculated by differential analysis between N_SON_ (or N_SRRM2_) and N_-priAB_. In Method 2, N_SON_ (or N_SRRM2_) was background subtracted by sequencing depth-normalized N_-priAB_ to remove any reads generated from antibody and/or pAG-RTase nonspecific binding, and then differential analysis was performed between the corrected N_SON_ (or N_SRRM2_) and N_nu-RNA_.

To demonstrate the robustness of our method to the choice of marker protein, we compared I_NSE_ values obtained from SON to those obtained from SRRM2. We found that using both analysis methods and in both HeLa and HepG2 cells, the I_NSE_ values obtained with the two marker proteins show a high degree of correlation (Figures S2B-C). Consistently, speckle-enriched transcripts (defined by I_NSE_ > 2 and adjusted p-value < 0.05) identified using SON and SRRM2 show a high degree of overlap (Figure 1C and S2D). The robustness of ARTR-seq to the choice of marker protein demonstrates that it could capture RNA in a defined 3D proximity, largely representing speckle-localized RNAs. Further supporting this idea, the well-known speckle-localized lncRNA *MALAT1* (Hutchinson et al., 2007), which was consistently identified with a high I_NSE_ with both marker proteins, showed similar read coverage patterns using both marker proteins (Figure 1D). While both marker proteins provided good results, we did notice that SON consistently identified more speckle-enriched transcripts with less variation than SRRM2 (Figures 1C and S2D-G). This is likely due to the higher speckle enrichment of SON (Dopie et al., 2020), and/or a better antibody quality. Therefore, in the rest of our analyses, we mainly focus on the I_NSE_ values determined using the SON antibody.

Speckle-enriched transcripts identified using analysis Method 1 and 2 also show a high degree of overlap (Figure S2E and F). To validate the sequencing data and compare the two analysis methods, we performed fluorescence in situ hybridization (FISH) imaging on RNA transcripts that were either identified as speckle-enriched or speckle-non-enriched (Figure 1E and Figure S3A). Theoretically, I_NSE_ should reflect the ratio between the RNA concentration inside nuclear speckles and that in the nucleoplasm. We calculated this ratio by measuring the fluorescence signals inside nuclear speckles and in the surrounding nucleoplasm (denoted by R_NS/NU_). Using both analysis methods, speckle-enriched transcripts demonstrate high R_NS/NU_, whereas the R_NS/NU_ values for speckle-non-enriched transcripts were close to 1 (Figure S3B). We did, however, find that I_NSE_ values calculated using Method 1 demonstrate a strong quantitative correlation with R_NS/NU_ values (Figure 1F). This could be because the N_-priAB_ control used in Method 1 is generated with a nearly identical protocol to that used to generate N_SON_ (or N_SRRM2_), leading to a more accurate differential analysis. We will therefore mainly focus on analysis method 1 in the rest of our analyses.

Finally, we compared our ARTR-seq results with the previously reported APEX-seq results (Figure S4) (Barutcu et al., 2022). I_NSE_ values demonstrate a positive correlation with the “Index 1” calculated from APEX-seq (Figure S4). This index provides ordinal (rank) information on speckle enrichment. However, quantitative enrichment information is not easily derivable from the APEX-seq result. This is due to the exogenous APEX2 fusion protein potentially perturbing gene expression, which necessitates additional controls to confidently identify enriched genes. By avoiding the use of exogenous fusion proteins, ARTR-seq can directly provide enrichment quantification. Additionally, the speckle marker proteins used in the APEX-seq study (SRSF1, SRSF7, RNPS1) are not as highly enriched in speckles as the scaffold protein SON and SRRM2 (Dopie et al., 2020).

In summary, ARTR-seq provides robust and quantitative information on the nuclear speckle transcriptome.

### Unspliced introns are enriched in nuclear speckles

We next compared the total number of reads mapping to an exon-intron boundary (EI, averaged over both splice sites) to the total number of reads mapping to an exon-exon junction (EE). Combining these two values provides a global estimate of the fraction of unspliced introns (calculated as EI/(EI+EE))). Under normal condition (no treatment, NT), we observed similarly low fraction of unspliced introns in ARTR-seq without antibody, in polyA RNA-seq, and in nuclear RNA-seq, whereas the fraction of unspliced introns in ARTR-seq with SON antibody is about 7 to 8-fold higher (Figure 2A). This comparison suggests that in general, speckle-localized transcripts contain more unspliced introns compared to nucleus-localized transcripts.

**Figure 2.**
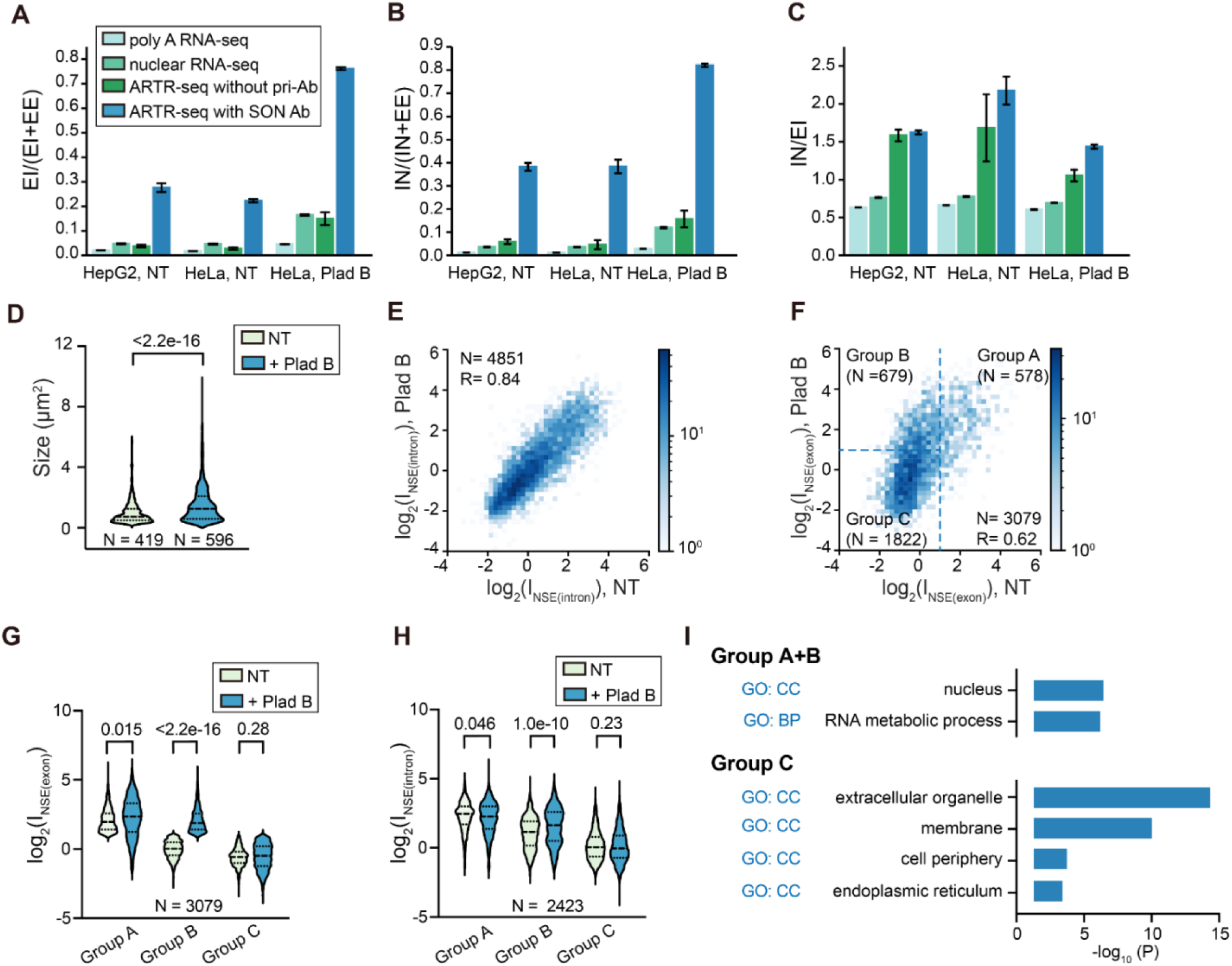
Transcripts exhibit varying nuclear speckle localization propensity and dynamics. (A) Fraction of unspliced introns (EI/(EI+EE)) calculated from the number of reads mapping to exon-intron boundary (EI) and the number of reads mapping to exon-exon junction (EE) in polyA RNA-seq, nuclear RNA-seq, ARTR-seq without antibody and ARTR-seq with SON antibody. (B) Alternative estimation of the fraction of unspliced introns (IN/(IN+EE)) calculated using reads mapping to intronic positions within 100 nucleotides of splice sites (IN) instead of EI. (C) Ratio of IN reads to EI reads in polyA RNA-seq, nuclear RNA-seq, ARTR-seq without antibody and ARTR-seq with SON antibody. The two replicates of RNA-seq were calculated individually. Each bar with error bars in (A)-(C) reports mean and standard deviation of the two replicates. (D) Violin plot showing the speckle size immunostained with SRRM2 antibody under NT Plad B treatment. “N” indicates the total number of speckles in each data set. (E-F) 2D histogram showing the correlation of I_NSE(intron)_ (E) or I_NSE(exon)_ (F) between NT and Plad B treatment conditions in HeLa cells. Genes were categorized into three Groups (Group A, B, C) and depicted in (F). (G-H) Violin plot comparing I_NSE(exon)_ (G) or I_NSE(intron)_ (H) values among Group A, B, and C genes under NT and Plad B treatment conditions. P-values calculated with unpaired t-test are reported above each violin plot. “N” reports the total number of genes in each comparison. (I) GO analysis for speckle-enriched Group A and Group B genes, and non-speckle-enriched Group C gene. The analysis is performed using g:Profiler (Raudvere et al., 2019), using the background consisting of all Group A, B and C genes. GO terms in biological processes (BP) and cellular compartment (CC) were identified.

To ensure the robustness of this calculation, we used the number of reads mapping to an intronic position within 100 nucleotides of the splice sites (IN, averaged over both splice sites) instead of EI to calculate the fraction of unspliced introns (calculated as IN/(IN+EE)), obtaining similar results (Figure 2B). We did notice, however, that there are consistently 1-2 fold more IN reads than EI reads in all ARTR-seq samples, including the one without antibody, but not in nuclear RNA-seq or polyA RNA-seq (Figure 2C). This is likely because in situ reverse transcription in crosslinked samples, as done in ARTR-seq, is sensitive to the presence of spliceosomal complexes around splice sites, slightly reducing the number of EI reads. A possible alternative explanation is that the number of IN reads is higher due to the presence of spliced intron lariats that have not yet been degraded or lariat intermediates. We find this possibility less likely since the effect was not observed in nuclear RNA-seq. In either case, this analysis suggests that the intronic read count reflects the abundance of unspliced introns and is not strongly affected by the presence of intron lariats or lariat intermediates.

### Splicing inhibition increases speckle localization of unspliced transcripts

It is suggested that most nascent transcripts are co-transcriptionally spliced (Brugiolo et al., 2013; Herzel et al., 2017). Due to the coupling between splicing and nuclear export (Reed and Hurt, 2002; Valencia et al., 2008; Viphakone et al., 2019), these rapidly spliced transcripts are subsequently exported. Any association of these transcripts with speckles during co-transcriptional splicing is therefore expected to be transient, and will not be frequently captured by ARTR-seq. It was observed that an increased amount of transcripts localize to speckles upon splicing inhibition (Kaida et al., 2007), suggesting that delaying splicing of certain transcripts increases their speckle enrichment. We therefore reasoned that inhibiting splicing may allow us to extend the time of speckle localization of these transcripts and better capture them in ARTR – seq.

We inhibited splicing by treating HeLa cells with 100 nM Pladienolide B (Plad B) for 4 h, and performed ARTR-seq. While Plad B cannot completely inhibit splicing for all endogenous genes (Kotake et al., 2007; Vigevani et al., 2017), we observed a 2-to 3.5-fold increase in the fraction of unspliced introns upon Plad B treatment across all experiments (Figure 2A). In particular, the fraction of unspliced introns in ARTR-seq with SON antibody is ∼76% upon Plad B treatment (Figure 2A), suggesting that the majority of speckle-localized transcripts are unspliced. These results confirm that splicing inhibition increases the speckle localization of unspliced transcripts, consistent with the observation of a ∼1.6-fold increase in median speckle size upon Plad B treatment (Figure 2D). However, in addition to extending the speckle localization of those transiently localized pre-mRNAs, we cannot rule out the possibility that additional transcripts become enriched in speckles due to splicing inhibition. It is also worth noting that the ratio between IN and EI decreases upon Plad B treatment (Figure 2C), supporting a decreased occupancy of spliceosomes at the splice sites.

### Transcripts demonstrate diverse dynamics in speckle localization

Motivated by the observed high ratio of intron reads in speckle-localized transcripts, we further calculated I_NSE_ values using either total exon reads (denoted as I_NSE(exon)_) or total intron reads (denoted as I_NSE(intron)_) mapped to each gene. We reasoned that I_NSE(exon)_ reflects speckle enrichment of “total transcripts”, i.e., including both unspliced transcripts and fully spliced but not yet exported mRNAs, whereas I_NSE(intron)_ mostly reflects speckle enrichment of unspliced transcripts. Therefore, a gene whose transcripts only transiently localize to speckles at the pre-mRNA state, but rapidly exit speckles upon splicing, is likely to demonstrate a high I_NSE(intron)_, but low I_NSE(exon)_. Moreover, under splicing inhibition, due to an increase in the fraction of the unspliced RNA in the total transcript, we expect such genes to show higher I_NSE(exon)_ compared to NT condition.

With this rationale, we compared enrichment values under NT and Plad B treatment. Surprisingly, we found a strong correlation between I_NSE(intron)_ values between NT and Plad B treatment (Figure 2E). Due to their total intronic length, fully transcribed with more unspliced transcripts contribute disproportionally to intronic read count, and are expected to dominate I_NSE(intron)_. The observed strong correlation therefore suggests that Plad B treatment overall has minimal impact on speckle localization propensity of these unspliced transcripts, but only increases their proportion among all transcripts. In contrast, we found that some genes exhibited large Plad B-dependent increase in I_NSE(exon)_ (Figure 2F). To facilitate further analysis, we separated genes into three groups based on I_NSE(exon)_ values (Figure 2F): Group A genes with log_2_(I_NSE(exon)_) > 1, i.e. being >2-fold enriched under NT; Group B genes with log_2_(I_NSE(exon)_) < 1 under NT, but log_2_(I_NSE(exon)_) > 1 upon Plad B treatment, i.e., showing Plad B-dependent speckle localization at total transcript level; and Group C genes with log_2_(I_NSE(exon)_) < 1 in both conditions.

Comparison of the I_NSE(exon)_ and I_NSE(intron)_ under NT and Plad B treatment conditions allowed us to infer the dynamics of transcript speckle localization under NT condition. Group A genes consistently demonstrate the highest I_NSE(exon)_ and I_NSE(intron)_ regardless of splicing inhibition. We interpret it as an indication that transcripts from Group A genes are stably localized to speckles already under NT condition. Group B transcripts, whose I_NSE(exon)_ is similar to that of Group C genes and much lower than that of Group A genes (Figure 2G), overall exhibit a significantly higher I_NSE(intron)_ compared to Group C genes under NT condition (Figure 2H). This feature of Group B genes supports our rationale above that pre-mRNAs from Group B genes transiently localize to speckles, and that the spliced transcripts exit speckles upon splicing, leading to the observed high I_NSE(intron)_ but low I_NSE(exon)_ under NT condition. Splicing inhibition increases the fraction of pre-mRNA in the total transcripts from Group B genes, and causes an increase in I_NSE(exon)_ to be closer to I_NSE(intron)_. Finally, Group C transcripts, which show low I_NSE(exon)_, also consistently show the lowest I_NSE(intron)_ regardless of Plad B treatment, suggesting that transcripts from this subset of genes are generally not localized to speckles throughout transcription or splicing. The insignificant change in I_NSE(exon)_ and I_NSE(intron)_ for Group C transcripts upon Plad B treatment indicates that for transcripts with low speckle localization propensity, Plad B treatment is unlikely to increase their speckle localization. In other words, the ratio of transcript concentration in speckle and nucleoplasm (reported by I_NSE_) is unlikely to change even though the overall nuclear portion of the transcripts increases upon splicing inhibition.

In summary, genes classified in Groups A, B, and C demonstrate different speckle localization propensity and dynamics: Group A transcripts are stably enriched in speckles; Group B transcripts are transiently enriched in speckles at pre-mRNA stage; and Group C transcripts are not speckle enriched (graphical abstract). GO analysis using total genes in all three groups as background revealed that speckle-enriched Group A and B genes are enriched in biological processes related to mRNA metabolism, and nucleus localization, whereas non-speckle-enriched Group C genes are enriched in cellular compartments of extracellular organelle, membrane, cell periphery, and endoplasmic reticulum (Figure 2I).

### Transcript speckle enrichment is positively correlated with localization of genes relative to speckles

Previous studies suggested that actively transcribed gene foci tend to be associated with nuclear speckles (Chen and Belmont, 2019; Galganski et al., 2017; Ilık and Aktaş, 2022). We therefore analyzed the correlation between transcript speckle enrichment and the proximity of the gene foci to nuclear speckles, as measured using the tyramide signal amplification sequencing (TSA-seq) (Y. Chen et al., 2018). We identified a positive correlation between TSA score and both I_NSE(intron)_ (Figure 3A) and I_NSE(exon)_ (Figure 3B). This suggests that transcripts transcribed from speckle-proximal gene foci tend to be more enriched in nuclear speckles, consistent with previous findings (Barutcu et al., 2022). The correlation with I_NSE(intron)_ is higher than that with I_NSE(exon)_; this supports the rationale that I_NSE(intron)_ reflects speckle enrichment of pre-mRNAs, which are mainly around transcription sites associated with gene foci, whereas I_NSE(exon)_ reflects localization of total transcripts either at the transcription sites or away from them. Finally, TSA scores from Group A and B genes are significantly higher than Group C genes (Figure 3C), supporting that transcripts that are either stably or transiently speckle enriched both have their DNA foci localized closer to speckles.

**Figure 3.**
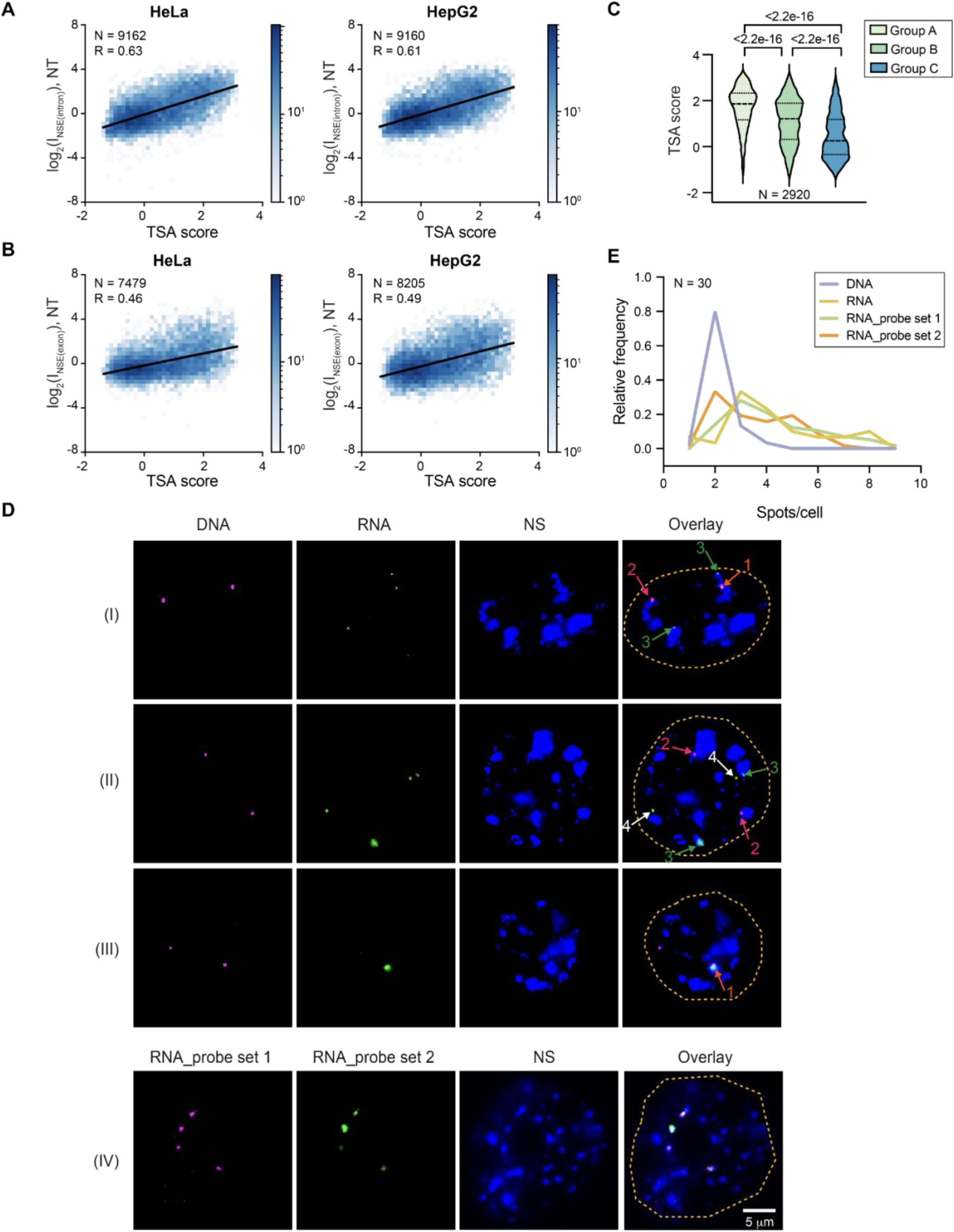
Correlation between RNA nuclear speckle enrichment and position of gene foci. (A-B) 2D histogram showing the correlation between I_NSE(intron)_ (A) or I_NSE(exon)_ (B) determined by ARTR-seq in HeLa cells or HepG2 cells and TSA score from TSA-seq in K562 cells. Genes with higher TSA scores are closer to speckles. (C) Violin plot comparing TSA scores for Group A, B, C genes. P-values calculated with unpaired t-tests are reported above each violin plot. “N” reports the total number of genes in the comparison. (D) Panel (I)-(III): Combined RNA FISH (labeled with CF568, green) and GOLD FISH (labeled with AF647, red) detection of speckle-enriched *LAMA5* transcripts and corresponding gene loci respectively. Arrows with different labels indicate different DNA/RNA localizations: (1) Speckle-associated colocalized RNA and DNA foci; (2) Speckle-associated DNA foci without co-localized RNA foci; (3) Speckle-associated RNA foci without co-localized DNA foci; (4) RNA foci that are not associated with a DNA focus nor localized to speckles. Panel (IV): RNA FISH detection of speckle-enriched *LAMA5* transcripts using two sets of probes labeled with AF647 and CF568. Colocalization of the AF647 and CF568 validates the specificity of transcript detection. (E) Histogram showing the distribution of the number of RNA foci and DNA foci per cell. “N” reports total number of cells included in the histogram, collected from 3 biological replicates.

We next investigated whether speckle-enriched transcripts are exclusively at active transcription sites. We imaged the intron regions of the speckle-enriched transcript *LAMA5* (from Group A) using RNA FISH together with its gene foci using the GOLD FISH method (Wang et al., 2021) (Figure 3D). On average, the number of RNA foci observed per cell is higher than that of DNA foci (Figure 3E). To exclude the possibility that the combination with GOLD FISH compromised the specificity of RNA FISH, we also performed RNA FISH alone using probes targeting different regions on *LAMA5* intron with two colors (Figure 3D). The two-color RNA FISH signals show ∼86% colocalization, and yield the same number of RNA foci per cell as in the combined DNA/RNA FISH imaging (Figure 3E), validating the specificity of RNA FISH.

Overall, 67±3% of *LAMA5* RNA foci are speckle-associated. However, only 40±9% of speckle-associated RNA foci have a DNA focus associated to the same nuclear speckle (Table S1), suggesting that RNA localization to speckle is not always associated with transcription sites. We further categorized RNA foci depending on their spatial relationship with the gene foci. The first kind of RNA foci are associated with an adjacent, nearly “touching”, DNA focus, and likely reflect co-transcriptionally accumulated RNAs at transcription sites. The second kind of RNA foci are physically separated from the DNA focus, and likely reflect post-transcriptionally accumulated RNAs. The DNA-associated RNA foci are nearly all speckle-associated (89±11%), whereas 60±1% of the DNA-detached RNA foci are speckle-associated (Table S1). In summary, co-staining of speckle-enriched RNA with the DNA foci supports that speckle localization can be both co-transcriptional and post-transcriptional.

### Transcript speckle enrichment is weakly correlated with RNA abundance

We next wondered whether transcript speckle enrichment is correlated with the expression level or the abundance of the transcript. We compared I_NSE(exon)_ and I_NSE(intron)_ values with gene expression levels measured by polyA RNA-seq, nuclear RNA abundance estimated by nuclear RNA-seq, as well as nascent transcript abundance measured by global run-on sequencing (GRO-seq) (Andersson et al., 2014). We observed an insignificant correlation between I_NSE(exon)_ and TPM values from polyA RNA-seq, nuclear RNA-seq, or GRO-seq, but weak correlations between I_NSE(intron)_ and TPM (transcript per million) values (Figure S5). These comparisons suggest that total transcript speckle enrichment is not correlated with gene expression level or transcription activity. However, speckle enrichment of pre-mRNAs is weakly correlated with gene expression level or transcription activity. Such correlations are also consistent with the previous claim that being closely associated with speckles may positively impact transcription (Chen and Belmont, 2019). The loss of correlation between I_NSE(exon)_ and TPM values from polyA RNA-seq, nuclear RNA-seq or GRO-seq, on the other hand, indicates that post-transcriptional localization of RNA is likely to be decoupled from transcript level or transcription activity.

### Transcript speckle enrichment is related to splicing kinetics and efficiency

We next analyzed whether speckle-enrichment of transcripts is related to splicing kinetics. We used data from a recent study using co-transcriptional lariat sequencing (CoLa-seq) (Zeng et al., 2022). By mapping intronic branch points, CoLa-seq reveals when an intron gets spliced relative to its adjacent introns. Specifically, in-order splicing (fast) represents the excision of an intron before transcription or splicing of the downstream intron; out-of-order splicing (slow) represents the excision of an intron after transcription and splicing of one or more downstream introns; and concurrent splicing (intermediate) reflects the excision of an intron around the same time as the downstream intron (Zeng et al., 2022). We calculated I_NSE_ values for individual introns following the same analysis used to calculate I_NSE_ at the transcript level, and then compared these values to CoLa-seq data. We found that out-of-order and concurrently spliced introns have significantly higher I_NSE_ values compared to in-order spliced introns (Figure 4A). This suggests that the presence of slowly spliced introns correlates with high transcript speckle enrichment. In addition, Group A genes contain significantly more introns with a small in-order splicing fraction, followed by Group B and Group C genes, suggesting that Group A genes are most enriched in slower spliced introns (Figure 4B).

**Figure 4.**
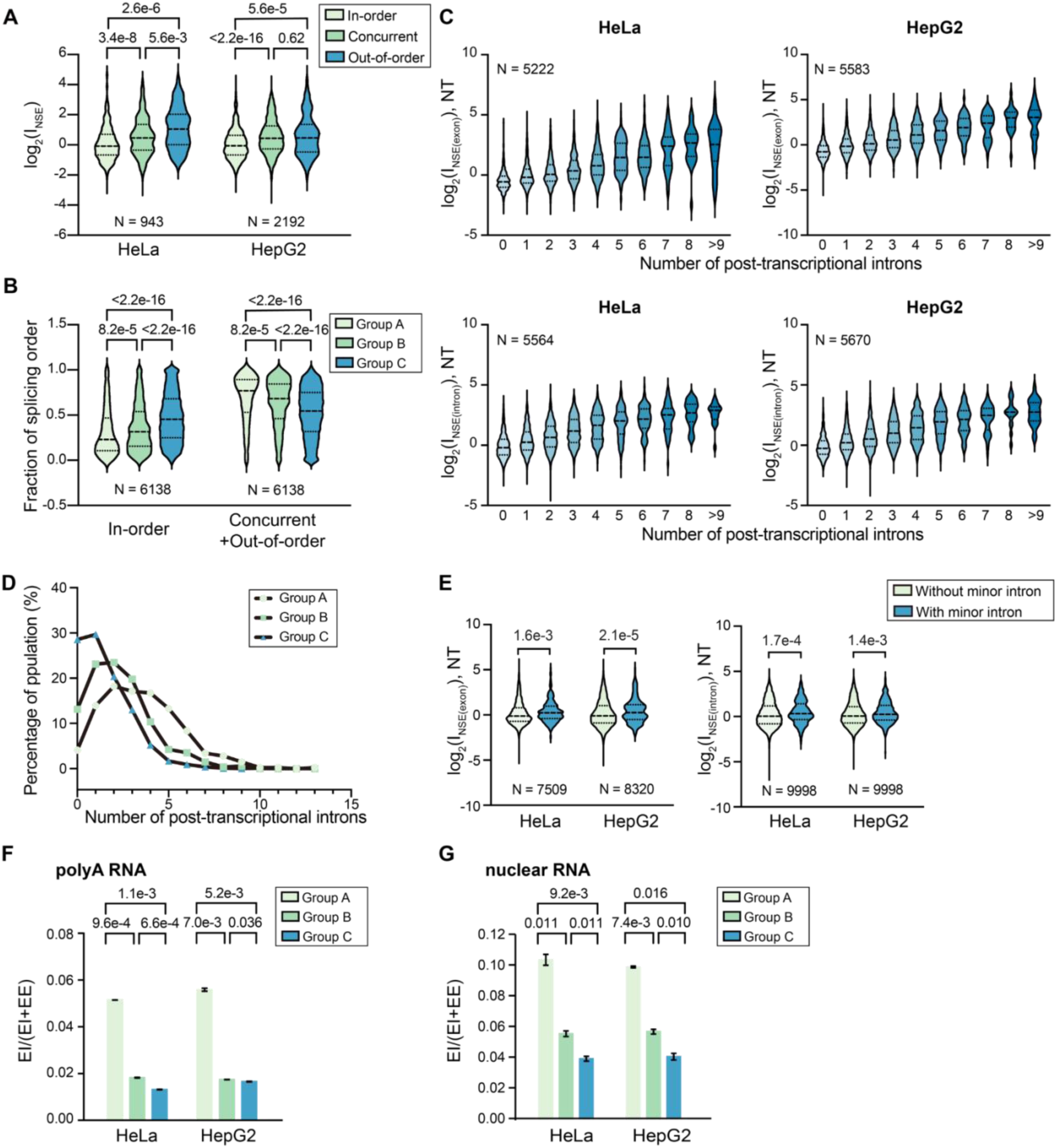
Transcript speckle enrichment is associated with splicing kinetics and efficiency. (A) Violin plot comparing I_NSE_ values of in-order spliced introns, concurrently spliced introns, and out-of-order spliced introns, as classified by CoLa-seq (Zeng et al., 2022). (B) Fraction of in-order spliced introns and non-in-order-spliced introns (concurrently spliced introns and out-of-order spliced introns) in Group A, B and C genes. (C) Violin plot comparing I_NSE(exon)_ or I_NSE(intron)_ values of transcripts containing different numbers of post-transcriptionally spliced introns, as characterized by nanopore RNA-seq in HeLa cells and HepG2 cells (Choquet et al., 2023). (D) Histogram showing the distribution of post-transcriptionally spliced intron number for Group A, B and C genes. (E) Violin plot comparing I_NSE(exon)_ or I_NSE(intron)_ values of transcripts containing minor splice sites and those without (Olthof et al., 2019). In (A)-(E), p-values calculated with unpaired t-tests are reported above each violin plot. “N” reports the total number of introns or genes in each comparison. (F-G) Fraction of unspliced introns (EI/(EI+EE)) for Group A, B and C genes under NT conditions at the polyA (F) and nuclear RNA level (G). The two RNA-seq replicates were calculated individually. Each bar with error bars in (F)-(G) reports mean and standard deviation of the two replicates. P-values calculated with unpaired t-tests.

To further investigate the effect of splicing kinetics on speckle localization, we compared I_NSE(exon)_ and I_NSE(intron)_ values of transcripts containing different numbers of post-transcriptional spliced introns, as characterized by a recent study using nanopore RNA-seq (Choquet et al., 2023). We found that on average, a transcript’s speckle enrichment increases with the number of post-transcriptionally spliced introns (Figure 4C). In addition, Group A genes contain significantly more post-transcriptionally spliced introns, followed by Group B and Group C genes, suggesting that Group A genes have a higher contribution from post-transcriptional splicing (Figure 4D).

Moreover, introns that use minor splice sites are known to be spliced slower (Patel et al., 2002, Zeng et al., 2022). We found that transcripts from genes containing introns utilizing minor splice sites (Olthof et al., 2019) are more enriched in speckles (Figure 4E). To summarize, these results further support that speckle enrichment is associated with transcripts containing slower spliced and post-transcriptionally spliced introns under NT condition, supporting the previous implication that nuclear speckles are involved in post-transcriptional splicing. In addition, for the two speckle-enriched gene groups, Group A genes demonstrate a higher post-transcriptional splicing propensity than Group B genes.

Finally, we compared the fraction of unspliced introns for Group A, B and C genes under NT condition. Group A genes consistently demonstrate a 2.8-3.9 fold higher value compared to Group B and C genes at the polyA RNA level (Figure 4F), and a 1.7-2.7 fold higher value compared to Group B and C genes at the nuclear RNA level (Figure 4G). This observation is consistent with the above analyses and supports that Group A genes undergo slower splicing and have a higher post-transcriptional splicing propensity than Group B and C genes.

### Nuclear speckles facilitate splicing of speckle-enriched transcripts

To test whether nuclear speckles facilitate splicing of speckle-enriched transcripts, we randomly chose a few genes from Groups A, B and C (Figures 5A). For each of these selected genes, we then picked introns that are either inefficiently spliced (showing intronic reads in polyA RNA-seq or nuclear RNA-seq), or efficiently spliced (not showing intronic reads) (Figures 5B and S6A). Using primers flanking the selected introns, we performed reverse transcription polymerase chain reaction (RT-PCR) analysis on cells that underwent either mock treatment (using control siRNA) or speckle disruption by double siRNA knockdown of the scaffold proteins SON and SRRM2 (Figures 5C-F). Upon double knockdown, immunostaining revealed a ∼70% decrease in SRRM2 signal (Figure 5E) and the average number of speckles per cell dropped from 22.5 to 2.2 (Figure 5F), indicating efficient speckle disruption.

**Figure 5.**
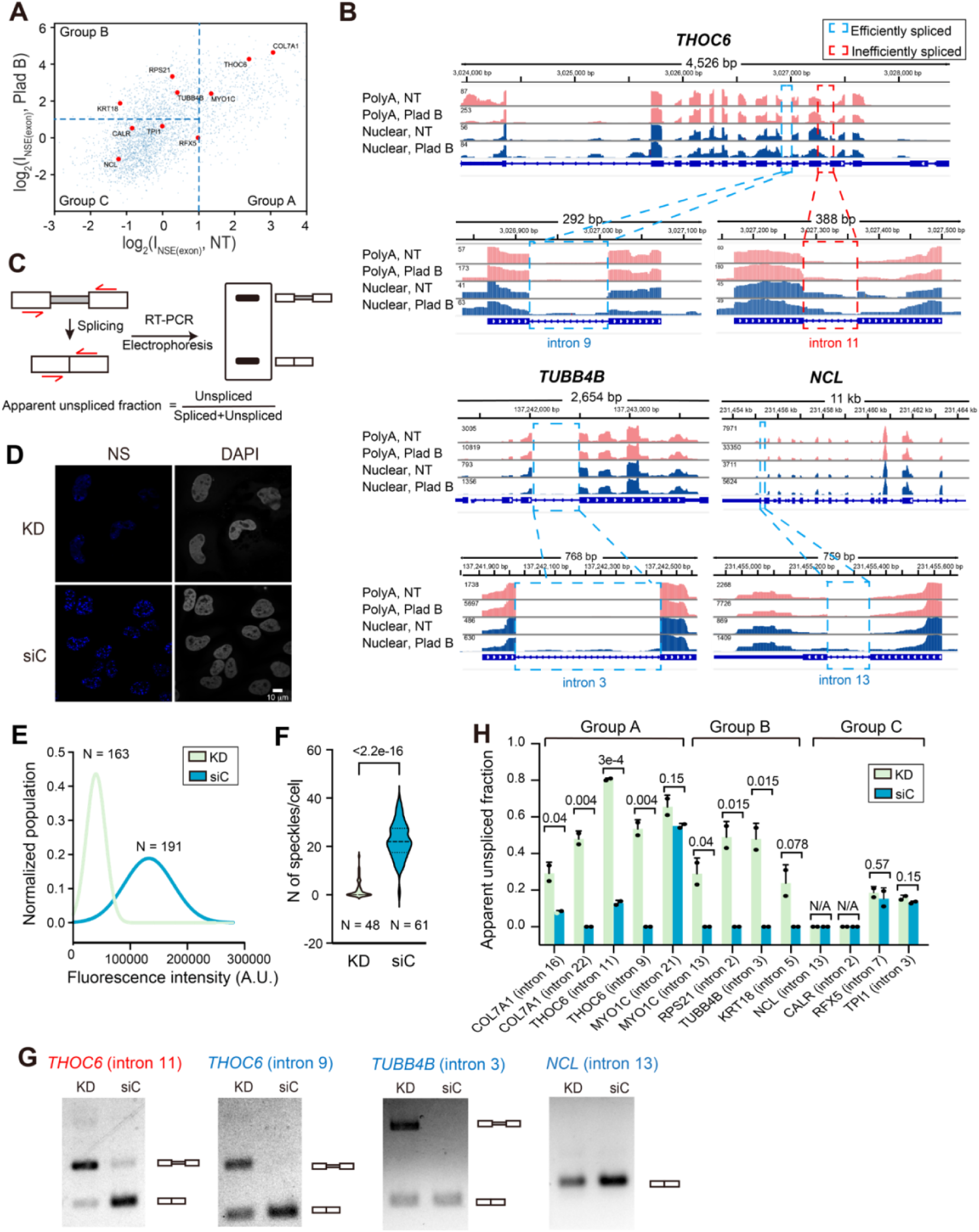
Nuclear speckles facilitate splicing of speckle-enriched transcripts. (A) Scatter plot showing randomly selected genes from Group A, B and C genes, and corresponding I_NSE(exon)_ under NT and Plad B treatment conditions in HeLa cells. (B) Genome tracks showing polyA RNA-seq (pink) and nuclear RNA-seq (blue) under NT and Plad B treatment conditions for selected genes: *THOC6* in Group A gene, *TUBB4B* in Group B gene, and *NCL* in Group C gene. Selected efficiently spliced or inefficiently spliced introns for RT-PCR assay are highlighted in cyan and red boxes respectively. Genome tracks of other selected genes for RT-PCR assays are shown in Figure S8. (C) Schematic description of the RT-PCR assay. After reverse transcription of extracted total RNA, primers located on two adjacent exons of selected introns were used for amplification and the PCR products were analyzed by electrophoresis. (D) Representative immunofluorescence images showing nuclear speckles upon *SON*/*SRRM2* double knockdown (KD) and treated with control siRNA (siC). Nuclear speckles were stained with AF488 labeled antibody against SRRM2 antibody (blue); and nuclei were stained with DAPI (grey). Scale bar: 10 μm. (E) Histogram of SRRM2 immunofluorescence intensity distribution of cells with double knockdown (KD) or treatment with control siRNA (siC). (F) Violin plot showing the number of speckles per cell for KD and siC treatment. Total number of cells included in each data set is indicated by “N” in (E) and (F). P-values calculated with unpaired t-tests are reported above each violin plot. (G) Representative electrophoresis analysis of RT-PCR products from *THOC6*, *TUBB4B* and *NCL* upon KD and siC treatment. Gels of other selected genes for RT-PCR assays are shown in Figure S8. Gels were imaged with Chemidoc Imaging System. (H) Apparent unspliced fractions of selected introns were calculated by ratios of the intensity of the unspliced band and the sum of the unspliced band and spliced band. The intensity of bands was quantified using Fiji. Error bars report standard deviation from two biological replicates.

Under mock treatment, the RT-PCR assay confirmed the RNA-seq data, showing that introns containing mapped reads are inefficiently spliced to various extents, whereas the rest of the introns are nearly fully spliced (Figures 5G-H and S6B). Interestingly, *SON*/*SRRM2* double knockdown impacted the removal of all tested introns in Group A and B transcripts but not in Group C transcripts (Figures 5G-H and S6B). These results suggest that nuclear speckles facilitate the splicing of transcripts in Groups A and B but not of those in Group C. This is consistent with the observation that Group C transcripts are not speckle-enriched (Figure 2F). In addition, the effect of speckle disruption on splicing of Group B transcripts further supports the idea that pre-mRNAs of Group B transcripts have intrinsic speckle localization propensity independently of splicing inhibition (Figure 2H) and utilize speckles for facilitating splicing even under no treatment condition. In contrast, pre-mRNA that only associates with speckles due to splicing inhibition is unlikely to be affected by our knockdown.

Collectively, these results suggest that nuclear speckles do not enhance splicing of all genes, but rather only of a subset of speckle-enriched transcripts. These results also suggest that speckle-facilitated splicing may occur both co-transcriptionally (for Group B transcripts, which are more transiently enriched in speckles at the pre-mRNA stage co-transcriptionally) and post-transcriptionally (for Group A genes, which demonstrate features correlated with post-transcriptional localization).

### A regression model predicts RNA sequence features associated with speckle enrichment

We next sought to identify cis-factors that contribute to nuclear speckle enrichment. Based on the above analyses, we reasoned that I_NSE(exon)_ under NT reflects stably speckle-enriched total transcripts, whereas I_NSE(intron)_ under Plad B treatment can best reflect speckle-enriched pre-mRNAs. Therefore, we focused on analyzing I_NSE(exon)_ under NT and I_NSE(intron)_ under Plad B treatment to specifically dissect features associated with stably speckle-enriched total transcripts and transiently speckle-enriched pre-mRNA respectively. We found that both I_NSE(exon)_ and I_NSE(intron)_ values demonstrate a consistent positive dependence on intronic and exonic GC content, and a negative dependence on the average intron length and total gene length, while the dependence on exon length is less obvious (Figure S7A-C).

To obtain a more detailed understanding of the relevant RNA sequence features, we used a regression model (generalized additive model) and fit it to the measured I_NSE_ values. The choice of input features to the model was based on the dependencies observed above and the correlation with splicing kinetics. Specifically, we included a gene’s GC content and its mean intron length as input features to the model. Since speckle enrichment demonstrates a similar correlation with intronic and exonic GC content (Figure S7), we did not separate these two in the regression model. In addition to these two gene-level input features, we also included several splicing-related features for each internal exon. These include the exon length, the strength of its flanking acceptor (3’) and donor (5’) splice sites, as determined using MaxEnt (Yeo and Burge, 2004), and a machine learning (ML) score of the exon sequence and of the flanking intronic sequences. This ML score is computed using a model trained on splicing assay data (Liao et al., 2023). It is high for sequences that are recognized as exons (such as those enriched in binding sites of SR family proteins) and low for sequences that are recognized as introns (such as those enriched in binding sites of hnRNP family proteins). Each of the latter four features (3’ splice site strength, 5’ splice site strength, exonic ML score, and intronic ML score) is quantile-binned into one of three bins, allowing to categorize each exon into one of 3^4^=81 possible combinations. To summarize, the input to the model consists of the gene’s GC content, its mean intron length, and a list of its internal exon features, each described by its length and a categorical value corresponding to its splice site strengths and ML scores. To arrive at its prediction, the model scores each of these separately, and outputs the total score. Using these features, the regression model achieved an excellent fit to the measured enrichment values (Figure 6).

**Figure 6.**
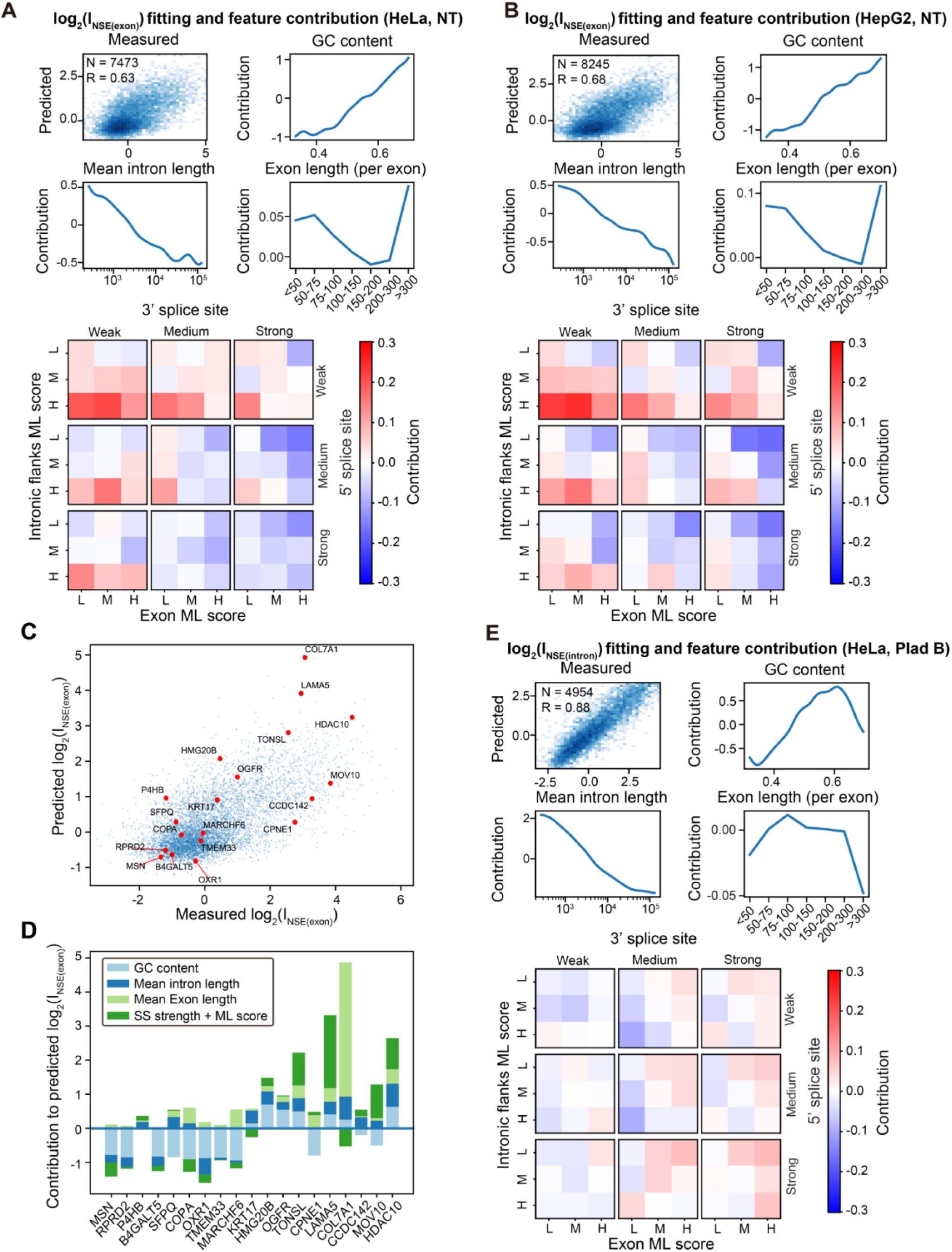
Regression model predicts RNA sequence features associated with speckle enrichment. Input parameters and other related details of the regression model are described in the main text and Methods. (A)-(B) The exon-centric regression model reveals contributions from GC content, mean intron length, individual exon length, and a combination of splice site strength, exonic ML score and flanking intronic ML score to the transcript speckle enrichment I_NSE(exon)_ values under NT in HeLa cells (A), under NT in HepG2 cells (B). (C) Predicted I_NSE(exon)_ values using the regression model on randomly selected is consistent genes are consistent with the measured I_NSE(exon)_ values from ARTR-seq. (C) The relative contribution from each parameter on selected genes. (E) The exon-centric regression model reveals contributions from GC content, mean intron length, individual exon length, and a combination of splice site strength, exonic ML score and flanking intronic ML score to the transcript speckle enrichment I_NSE(intron)_ values under Plad B treatment in HeLa cells.

### Speckle-enriched RNAs under NT condition demonstrate sequence features associated with inefficient splicing

We next interpreted the regression model in order to understand how the various features correlate with transcript speckle enrichment under NT condition (Figure 6A-B). Consistent with our previous correlation analysis, when analyzing I_NSE(exon)_ under NT, we found that high GC content and low mean intron length contribute significantly to speckle enrichment. Interestingly, the model revealed that short exons (<75nt) contribute to speckle enrichment, possibly related to the fact that they do not splice efficiently (Arias et al., 2015; Dominski and Kole, 1991). In addition, exons with a combination of weak splice sites and a high ML score for the flanking intronic sequences (suggesting those intronic sequences are not well-defined) are positively correlated with speckle enrichment. A mild contribution from a low ML score for the exon sequence (suggesting an exon that is not well-defined) was also observed. The same effects were consistently revealed in both HeLa and HepG2 cells (Figure 6A-B).

We further tested an alternative “intron-centric” regression model (Figure S8A-B), in which internal introns are categorized instead of internal exons. Specifically, for each internal intron, we used the strength of its flanking 5’ and 3’ splice sites, its ML score, and the ML score of the flanking upstream and downstream exons. These features were binned and combined as above, allowing us to label each internal intron with one of 81 possible combinations. The remaining features (GC content, mean intron length, and exon lengths) were kept the same. The sequence features identified by this model are largely consistent with the previous exon-centric model. Namely, high GC content, short mean intron length, short exon lengths, and a combination of weak splice sites and high intronic ML scores are all positively correlated with speckle enrichment, though the dependence on exonic ML score was not obvious in the intron-centric model.

To demonstrate the regression model’s prediction process, we randomly selected several genes and used the model to predict their speckle enrichment from sequence features. In agreement with the model’s good fit, the predicted values are well correlated with the I_NSE(exon)_ values experimentally measured by ARTR-seq (Figure 6C). Interestingly, the predictions are not dominated by any one feature; instead, each sequence feature (GC content, mean intron length, exon lengths, and splice site strengths with ML scores) can be the major contributing factor in a transcript-dependent manner (Figure 6D). A similar transcript-dependent feature contribution was also observed for predicting splicing timing (Zeng et al., 2022).

In summary, the regression analysis on I_NSE(exon)_ values under NT reveals features that distinguish Group A transcripts from Group B and C transcripts under normal condition. Features including high GC content, short introns and exons, and a combination of weak splice sites with high intronic ML score all contribute to speckle enrichment. We refer to these features collectively as Type I features. Interestingly, similar features (namely, high intronic GC content, short intron length, weak splice sites, and enrichment of intronic SR protein binding motifs) were previously reported for retained introns (Braunschweig et al., 2014; Middleton et al., 2017; Monteuuis et al., 2019; Schmitz et al., 2017). These results suggest that transcripts that are difficult to splice fully (due to the presence of weak splice sites or suboptimal cis-factors within exons or introns) are preferentially enriched in nuclear speckles, likely post-transcriptionally. These results are also in line with the hypothesis that nuclear speckles participate in post-transcriptional splicing.

### Speckle-localized pre-mRNAs demonstrate sequence features associated with efficient splicing

Performing the same analysis on I_NSE(intron)_ under NT values revealed similar dependence on GC content and intron length as in the previous I_NSE(exon)_ analysis. However, features associated with splicing are strongly diminished (Figure S8C). Since I_NSE(intron)_ reflects pre-mRNA speckle enrichment, the disappearance of splicing-related features indicates that speckle-enriched pre-mRNAs may not exhibit the same splicing-related features as speckle-enriched total transcripts.

Since splicing inhibition increases the contribution of pre-mRNA, we repeated the same regression analysis on I_NSE(intron)_ under Plad B treatment (Figure 6E). GC content and intron length dependence were robustly identified. In contrast, this analysis revealed different splicing-related features, which are largely opposite of those identified in the I_NSE(exon)_ analysis (Figures 6A-B). Specifically, speckle-enriched genes exhibit a moderate preference for strong 5’ and 3’ splice sites in combination with a high exonic ML score. We collectively refer to these features together with a preference of high GC content and short intron length as Type II features, which are more associated with speckle-localized pre-mRNAs. As strong splice sites and strong exonic ML score are features associated with efficient splicing (Liao et al., 2023), this correlation indicates that these pre-mRNA transcripts undergo efficient splicing.

Independent analysis of GC content, intron length, and splicing-related features in Group A, B, and C transcripts further confirmed that speckle-localized Group A and B transcripts have distinct features (Figure S7D). Group A transcripts have the highest GC content, shortest average intron length, and a preference for a combination of weak spice sites and high intronic ML score. Group B transcripts have an intermediate GC content, an intermediate average intron length, and a preference for the combination of strong splice sites with high exonic ML score. Finally, Group C transcripts have the lowest GC content, longest average intron length, and a preference for low intronic and exonic ML score independent of splice site strength (graphical abstract).

### Nuclear speckles serve as a stress response hub

Membraneless organelles play important roles in stress response (Biamonti et al., 2010; Protter et al., 2016; Riggs et al., 2020). For example, upon stress, translationally-paused mRNA can be temporarily sequestered to cytoplasmic stress granules (Protter et al., 2016). Our results demonstrate that nuclear speckles accommodate transcripts with inefficiently spliced introns (Group A genes). Since intron retention is known to regulate gene expression through diverse mechanisms (Braunschweig et al., 2014; Monteuuis et al., 2019) including nuclear detention (Boutz et al., 2015; Monteuuis et al., 2019; Morgan et al., 2019; Parenteau et al., 2019), we wondered whether cells utilize speckles to respond to stress.

To address this question, we used heat shock as an example. We stressed the cells at 43°C for 2 h and performed polyA RNA-seq. Consistently with previous results using mouse fibroblasts (Shalgi et al., 2014), we observed an overall upregulation of intron retention in polyA RNA-seq upon heat shock as identified by IRFinder (Middleton et al., 2017), with >4-fold more upregulated than downregulated intron retention events (Figure 7A). We checked whether transcripts containing heat shock-induced retained introns are functionally relevant under this specific stress case. We found that none of the classic heat-responsive heat shock proteins (Kampinga et al., 2009; Mahat et al., 2016), including those belonging to the HSPA (HSP70), HSPB (small HSP), HSPC (HSP90), HSPH (HSP110) and DNAJA/DNAJB (HSP40) families, exhibits heat-induced intron retention increase (with *HSPE1* being the only exception). Consistently, GO enrichment analysis of genes containing introns demonstrating >15% increased retention upon heat shock (ΔIR_>15%_) did not identify any heat response-related term, no matter whether the analysis background was chosen as the whole genome or as all expressed genes. These results suggest that intron retention may be employed to negatively regulate the expression of non-heat stress-related genes, providing a survival benefit.

**Figure 7.**
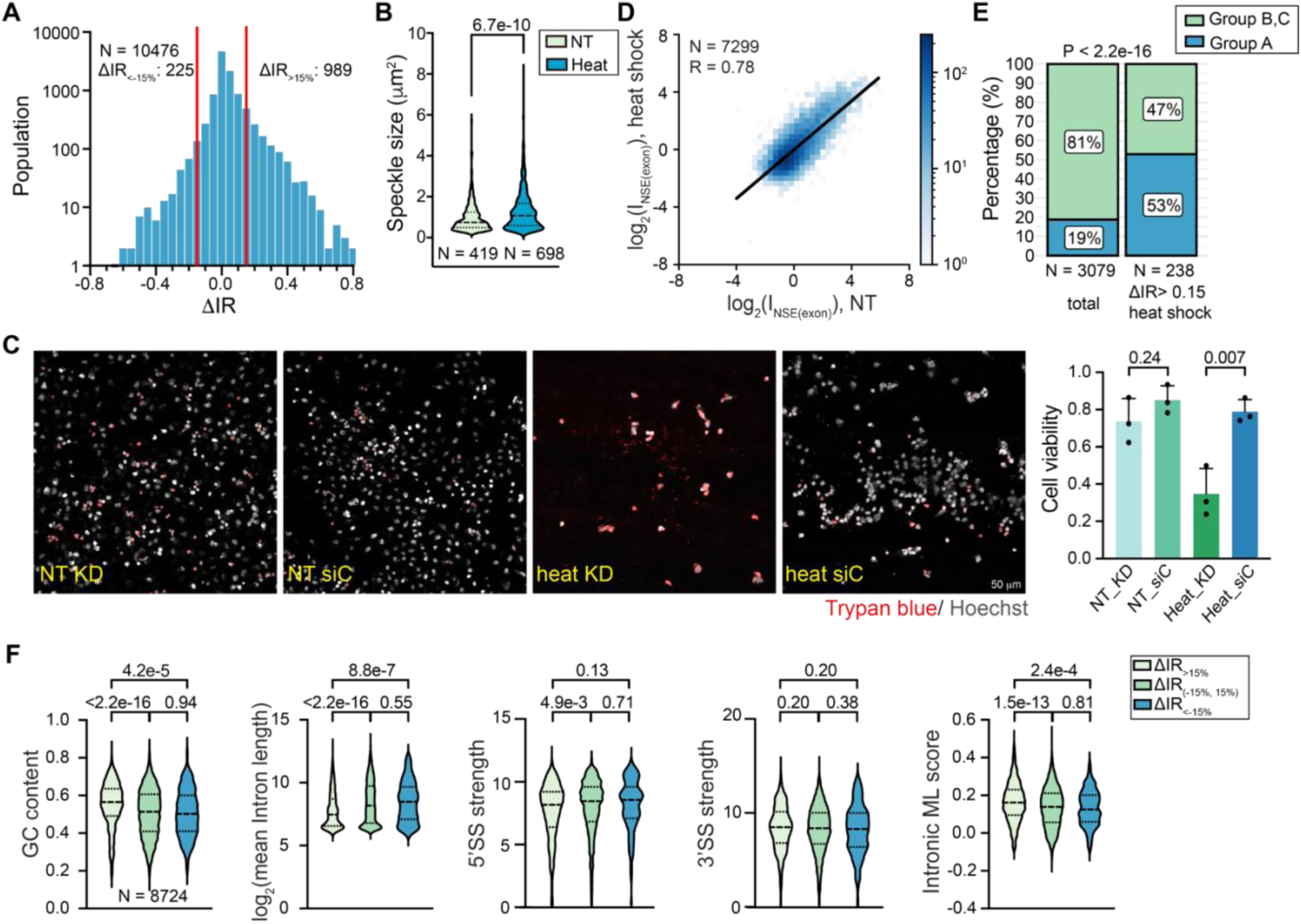
Functional implication of nuclear speckle under heat shock. (A) IRFinder analysis showing heat shock-induced upregulation of intron retention. The number of intron retention events with more than 15% increase (ΔIR_>15%_) or decrease (ΔIR_<-15%_) are labeled. (B) Violin plot showing the speckle size change up heat shock compared to NT. “N” indicates the total number of speckles in each data set. (C) Viability of HeLa cells with nuclear speckle disrupted using SON/SRRM2 double knockdown (KD) or cells treated with control siRNA (siC) upon heat shock stress or NT. Hoechst staining reflected the whole cell population (with cell number denoted as N_Total_), whereas Trypan blue stained the dead cell (with number denoted as N_Dead_). Cell viability was calculated by 1-N_Dead_/N_Total_. P-values calculated with unpaired t-test are reported above each violin plot and box plot. Error bars report standard deviation from 3 biological replicates (in black dots). (D) 2D histogram showing the correlation between I_NSE(exon)_ values under heat shock and NT. (E) Percentage of Group A genes and Group B and C genes without and with taking the subset of genes containing ΔIR_>15%_ introns. P-value: Fisher’s exact test. (F) Violin plot showing the Type I sequence feature associated with three groups of introns (ΔIR_>15%_, ΔIR_(–15%, 15%)_, ΔIR_<-15%_). The GC content, intron length, splice site strength and intronic ML score are compared for three groups of introns.

We next sought to analyze the relationship between heat shock-induced intron retention and transcript speckle enrichment. First, we observed a significant increase in speckle size upon heat shock (Figure 7B). We also found that cells with *SON*/*SRRM2* double knockdown demonstrate reduced viability upon heat shock compared to the ones treated with the control siRNA (Figure 7C), hinting at a possible role of speckles in heat shock response. To examine this possibility, we next performed ARTR-seq on the stressed cells. Interestingly, comparing I_NSE(exon)_ values under NT and under heat shock revealed a strong correlation (Figure 7D), unlike what was observed for the splicing inhibition perturbation (Figure 2F). Consistently, when applying the regression analysis on I_NSE(exon)_ values under heat shock, we found the same Type I sequence features as we identified under unstressed condition (Figures S8D). A previous study illustrated that transcripts with heat shock-induced intron retention are retained in the nucleus (Shalgi et al., 2014). Our results further suggest that increased abundance of transcripts with retained introns leads to an increased abundance of speckle-localized transcripts; however, the ratio between transcript concentration in speckles and in the nucleoplasm (measured by I_NSE(exon)_) for a particular gene remains largely unchanged. In summary, while heat shock does not significantly affect transcript speckle enrichment, the increase in speckle size does suggest that speckles play a role in storing and possibly processing transcripts with heat shock-induced intron retention.

To explore this possibility, we compared intron retention changes to I_NSE(exon)_ values. We found that 53% of classified ΔIR_>15%_ genes are in Group A, even though Group A genes only occupy 19% of all classified genes (Figure 2F and 7E). To further support this correlation, we inspected the sequence features exhibited by heat shock-induced retained introns. We classified retained introns into three groups: ΔIR_>15%_, for introns demonstrating >15% increased retention upon heat shock, ΔIR_(–15%, 15%)_, for those with insignificant retention level change, and ΔIR_<-15%_ for those demonstrating >15% decreased retention. We found that ΔIR_>15%_ introns exhibit stronger Type I features compared to the other two groups. Specifically, such introns exhibited significantly higher GC content, shorter intron length, weaker 5’ splice site, and stronger intronic ML score (Figure 7F). In summary, Group A genes and introns sharing the Type I features demonstrate increased intron retention levels.

Collectively, these analyses suggest that speckle-enriched genes tend to be more sensitive to perturbation at the splicing level, pointing to nuclear speckles serving as a hub for stress-induced intron retention upregulation.

## DISCUSSION

In this work, we broadened the applicability of our recently developed ARTR-seq method to transcriptomically map nuclear speckles. While ARTR-seq was originally developed as a method to identify direct protein binding sites, given the high density of speckle-localized RNA, it is also robust in capturing transcripts in the vicinity of the antibody anchoring point. The captured RNAs might be direct or indirect binding targets of SON and SRRM. Distinguishing the two possibilities is beyond the focus of the current work and our results are valid in either case. Compared to a related transcriptomic analysis on nuclear speckles using APEX-seq (Barutcu et al., 2022), our study provides an alternative approach with several advantages. First, our approach can better preserve the integrity of nuclear speckles, by directly targeting endogenous speckle marker proteins, and by avoiding potential cell stresses caused during sample treatment, such as using hydrogen peroxide. Second, compared to using SRSF1 and SRSF7 as speckle-targeting proteins (Barutcu et al., 2022), we target the most speckle-enriched scaffold proteins SON and SRRM2 (Dopie et al., 2020), thereby increasing targeting specificity to nuclear speckles. Third, the method is very flexible, and can be readily adapted to other marker proteins of interest without requiring the generation of fused proteins. Finally, in situ reverse transcription avoids the use of diffusive radicals, potentially increasing the localization accuracy for studying membraneless organelles compared to APEX-seq.

Comparing enrichment under NT and under Plad B treatment allowed us to classify genes into three groups with different speckle localization dynamics (graphical abstract). Transcripts from Group A genes are speckle-enriched at the pre-mRNA stage (high I_NSE(intron)_ under NT), likely co-transcriptionally, and remain enriched post-transcriptionally (high I_NSE(exon)_ under NT), presumably due to the presence of one or more slowly spliced or retained introns. Transcripts from Group B genes are also speckle-enriched at pre-mRNA stage (high I_NSE(intron)_ under NT), but exit speckles after splicing (low I_NSE(exon)_ under NT). Finally, transcripts from Group C genes are not enriched in speckles (low I_NSE(intron)_ and low I_NSE(exon)_ under NT). Importantly, disruption of nuclear speckles affects splicing efficiency of both Group A and B transcripts, but not Group C transcripts. Collectively, our data reveal that nuclear speckles facilitate both co– and post-transcriptionally splicing for a subset of genes.

Our data support the previous observation that not all actively transcribed genes are speckle-associated (Chen and Belmont, 2019), and reveal a tight interplay between genomic organization, RNA localization to nuclear speckle, and sequence features. Consistent with previous results from APEX-seq, we found that gene proximity to nuclear speckles is moderately correlated with total transcript speckle enrichment, and more strongly to the pre-mRNA speckle enrichment. Consistently, Group A and B gene foci are both closer to nuclear speckles compared to Group C genes. Co-staining of RNA transcripts with its gene foci supports that transcript localization to speckle can occur at the transcription site, likely co-transcriptionally, or away from transcription site, likely post-transcriptionally. Interestingly, we also found that some Group C genes, such as *CALR* and *TPI1*, have a high TSA score, suggesting that gene position alone cannot explain transcript localization to speckles. Using regression analysis, we further uncovered Type I and II sequence features that positively contribute to RNA localization to nuclear speckles, and found that having short GC-rich introns is associated with higher transcript speckle enrichment, both at the total transcript and at the pre-mRNA levels. Interestingly, genes containing such introns are organized in the interior region of the nucleus (Tammer et al., 2022). Therefore, the association of these features with Group A and B genes might be directly related to genome organization. Namely, it is possible that gene position impacts transcript speckle localization. Alternatively, cis-elements leading to transcript speckle localization facilitate the recruitment of speckles to the transcription site or the movement of gene foci toward nuclear speckles.

Our regression analysis also identified correlation between transcript speckle localization and splicing-related features, suggesting a model in which nuclear speckles coordinate splicing both co– and post-transcriptionally. Features such as weak splice sites in combination with intronic localized SR protein binding motifs (reflected by high intronic ML score) appear in Group A genes (whose total transcripts are speckle enriched), supporting the role of nuclear speckles as a processing site for slowly spliced or retained introns. In contrast, when considering speckle enrichment at pre-mRNA level, the regression analysis identified some splicing favored elements, such as strong splice sites in combination with exonic localized SR protein binding motifs (reflected by high exonic ML score), as being associated with higher pre-mRNA speckle enrichment. These correlations support the hypothesis that pre-mRNAs with Type II features are co-transcriptionally localized to speckles. Group B transcripts, which are globally more efficiently spliced than Group A transcripts, rapidly exit speckles upon splicing and are subsequently exported into cytoplasm. Group A transcripts with Type I features, which are more enriched in slowly spliced introns and globally contain a higher fraction of unspliced introns than Group B transcripts, are further retained and spliced in nuclear speckles post-transcriptionally.

While the exact mechanisms underlying the correlation between RNA cis-elements and speckle localization remain to be investigated, we hypothesize the following factors: (1) High GC content in the speckle-associated Group A and B transcripts may naturally have a higher propensity to partition into phase-separated domains (Jain and Vale, 2017; Wang and Xu, 2024). (2) Similarly, the presence of more SR protein binding motifs associated with Group A and B transcripts may contribute to speckle localization given the speckle enrichment of many SR proteins (Dopie et al., 2020). (3) Splicing promoting Type II features in Group B transcripts may facilitate spliceosome assembly on these pre-mRNAs, which increases their speckle localization given the known speckle-enrichment of spliceosomal components (Dopie et al., 2020; Fu and Ares, 2014; Galganski et al., 2017).

Our model of nuclear speckles participating in both co– and post-transcriptional splicing is consistent with earlier observations using imaging. For example, a post-transcriptionally spliced intron (intron 24 of *COL1A1*) and a retained intron (mutation-containing intron 26 of *COL1A1*) were observed to accumulate in nuclear speckles (Hall et al., 2006; Johnson et al., 2000). These imaging experiments also revealed intra-speckle positional differences between co– and post-transcriptionally spliced introns in *COL1A1*: co-transcriptionally spliced introns stay at the periphery of speckles with the gene foci outside speckles, whereas post-transcriptionally spliced introns and retained introns are distributed throughout the speckle. Therefore, it is possible that transcripts with Type II sequence features and inefficiently spliced transcripts with Type I sequence features may have different intra-speckle localization. However, further investigations are needed to clarify the functions of the speckle core and outer shell in coordinating co– and post-transcriptional splicing.

Repression of genes that are not directly needed for stress response can occur at the splicing level through intron retention (Boutz et al., 2015; Monteuuis et al., 2019; Morgan et al., 2019; Parenteau et al., 2019). Our results suggest that nuclear speckles serve as a hub for stress-regulated intron retention. Indeed, we found that under heat shock, a global increase in intron retention is correlated with an increased speckle size. Moreover, Group A genes (whose transcripts are speckle enriched) are highly over-represented in genes demonstrating heat shock-induced increase in intron retention. Consistently, introns retained under heat shock demonstrate Type I sequence features, which we found to be associated with speckle localization. These correlations suggest that Group A transcripts are more sensitive to heat stress-induced changes in splicing factors, likely due to the presence of Type I sequence features, demonstrate upregulated intron retention and are further accumulated to speckle. In other words, cells utilize the shared Type I features between retained introns and speckle-localized Group A transcripts as one way to negatively regulate gene expression at splicing level with speckles as a storage site for transcripts with retained introns. Moreover, the enrichment of spliceosomal components in speckles (Galganski et al., 2017; Ilık and Aktaş, 2022) may facilitate splicing upon recovery from stress. Further investigations are needed to fully elaborate the relationship between stress, intron retention and nuclear speckle localization.

### Limitations of the study

ARTR-seq tends to provide relatively short reads, preventing confident isoform-level analysis. It also produces uneven read coverage within transcripts, complicating attempts to identify intra-molecular speckle localization differences. These limitations can be partially attributed to the accessible range of RTase. While they do not affect our current analysis, further optimization of the method can potentially overcome them, allowing for a more detailed analysis of the interaction between transcription, splicing, and nuclear speckles. While Plad B is commonly used as splicing inhibitor, its effect is shown to have sequence-dependence (Kotake et al., 2007; Vigevani et al., 2017), which may cause basis in our analysis result on identifying RNA cis-elements that are contributed to speckle localization for pre-mRNAs. Finally, while we assume most pre-mRNA undergo splicing co-transcriptionally, and interpret I_NSE(intron)_ to mostly reflect co-transcriptionally spliced pre-mRNAs under no treatment conditions, current ARTR-seq experiments cannot distinguish co-versus post-transcriptional splicing. It is very likely that a fraction of the Group B pre-mRNAs, undergo rapid post-transcriptional splicing after transcription termination at nuclear speckles.

### Data and code availability

All the sequencing data generated in this study will be available upon acceptance of this manuscript. Previously published data are available under accession numbers GSE62046 (GRO-seq). This paper does not report any original code. Any additional information required to reanalyze the data reported in this paper is available from the lead contact upon request.

## Author contributions

Conceptualization: JF, JW, YX

Experiment: JW, YX, SP

Analysis: JW, YX, YL, LW, CJ, SL, OR, JF

Supervision: JF, OR, CH Writing: JF, OR, JW, YX

## ACKNOWLEDGEMENTS

We thank Dr. Jonathan P. Staley for sharing the CoLa-seq data, useful discussion and comments on the manuscript; Dr. Taekjip Ha, Dr. Yanbo Wang and Mr. Taylor Cottle for sharing GOLD FISH reagents and protocols; Dr. Susan E. Liao for comments on the manuscript; Mr. Wei Liu and Dr. Zunwu Zhou for managing Fei lab; Dr. Pieter Faber and the staff at the University of Chicago Genomics Facility for their support in high-throughput sequencing. This project was supported by the NIH Director’s New Innovator Award (1DP2GM128185-01) to JF, a Simons Investigator Award and NSF MCB-2226731 to OR, and an NSF MCB-2246530 to JF and OR, and NIH RM1 HG008935 to CH. CH is an investigator of the Howard Hughes Medical Institute.

CH is a scientific founder, a member of the scientific advisory board and equity holder of Aferna Green, Inc. and a scientific co-founder and equity holder of Accent Therapeutics.

## METHODS

### Cell culture and treatment

HeLa human cervical cancer cells and HepG2 human hepatocellular carcinoma cells were cultured in high-glucose Dulbecco’s modified Eagle medium (DMEM, Gibco) supplemented with 4.5 g/L glucose, 1 mM sodium pyruvate, 50 U/mL penicillin/streptomycin solution (Gibco) and 10% fetal bovine serum (FBS, Gibco). Mycoplasma contamination was regularly tested for both cell lines. For fluorescence imaging, HeLa cells were seeded at a density of 3 × 10^4^ cells in an 8-well imaging chamber (#1.5 cover glass, Cellvis) and grown overnight to 70-80% confluency. For HepG2 cells, chamber was coated with 100 μL of Matrigel matrix (Corning, 5 mg/mL) at 37°C for 1 h before seeding the cells.

For splicing inhibition experiment, cells were treated with Plad B (100 nM, Cayman Chemical) at 37°C for 4 h in DMEM medium. For heat shock, cells were incubated at 43°C for 2 h before following experiments.

### *SON* and *SRRM2* knockdown

Knockdown of *SON* and *SRRM2* in HeLa cells was performed using Lipofectamine RNAiMAX Transfection Reagent (Thermo Fisher Scientific). siRNAs were designed and purchased from IDT. The cells were sequentially transfected with siRNA (*SON*) and siRNA (*SRRM2*) with a 24 h interval between each transfection with a final concentration of 5 nM. The cells were also transfected with the same concentration of control siRNA twice as a negative control. The cells were subsequently incubated at 37°C for an additional 48 h before further experiments.

### PolyA RNA-seq and nuclear RNA-seq

#### Nuclei isolation

HeLa cells or HepG2 cells were collected by centrifugation at 500 g for 3 min and washed once with 1 mL DPBS. The cell pellet was resuspended in 200 μL ice-cold lysis buffer (10 mM Tris-HCl, pH = 7.5, 0.15% NP40, 150 mM NaCl), and incubated on ice for 5 min. Then the cell lysate was gently pipetted up over 500 μL chilled sucrose cushion (24% RNase-free sucrose in lysis buffer), and centrifuged at 15,000 g for 10 min at 4 °C. The pellet was collected as nuclei.

#### RNA extraction

Total RNA from nuclei or whole cells was purified with TRIzol reagent (Thermo Fisher Scientific) according to the manufacturer’s instructions. RiboMinus Eukaryote kit (Thermo Fisher Scientific) was used to remove rRNA from nuclei RNA. Dynabeads mRNA DIRECT™ kit (Thermo Fisher Scientific) was used to extract polyadenylated RNA (polyA RNA) from total RNA. The RNA concentration was measured by NanoDrop 8000 Spectrophotometer (Thermo Fisher Scientific).

#### RNA sequencing

RNA sequencing libraries of rRNA-depleted nuclear RNA or polyA RNA were prepared with SMARTer Stranded Total RNA-Seq Kit v2 (Takara) according to the manufacturer’s protocols. Sequencing was performed at the University of Chicago Genomics Facility on an Illumina NovaSeq 6000 platform in single-end mode with 100 bp.

### ARTR-seq

ARTR-seq was performed according to the previously published procedure (Xiao et al., 2024). Briefly, HeLa or HepG2 cells were fixed with 1.5% paraformaldehyde (PFA) for 10min at room temperature, quenched with 125 mM glycine, and permeabilized with 0.5% Triton X-100 on ice for 10 min. Samples were blocked with 1 mg/mL UltraPure BSA (Thermo Fisher Scientific), stained with SON or SRRM antibodies at room temperature for 1 h, and then stained with fluorophore-labeled secondary antibody at room temperature for 30 min. Samples were then incubated with pAG-RTase for an additional 30 min. A reverse transcription reaction mixture was prepared by mixing 2 μM adapter-RT primer, 0.05 mM biotin-16-dUTP (Jena Bioscience), 0.05 mM biotin-16-dCTP (Jena Bioscience), 0.05 mM dTTP (Thermo Fisher Scientific), 0.05 mM dCTP (Thermo Fisher Scientific), 0.1 mM dATP (Thermo Fisher Scientific), 0.1 mM dGTP (Thermo Fisher Scientific), 1 U/μL RNaseOUT (Thermo Fisher Scientific) in 50 μL buffer of DPBS supplemented with 3 mM MgCl_2_. In-situ reverse transcription was performed by adding RT reaction mixture to cells and incubating at 37 °C for 30 min, and then quenched by adding 20 mM EDTA and 10 mM EGTA. To check the success of in situ reverse transcription, cells were stained with biotin monoclonal antibody (BK-1/39) conjugated with Alexa Fluor 488 (AF488, Thermo Fisher Scientific), and imaged by Leica SP8 laser confocal microscope. The fluorescence intensity distribution on a line was quantified by Fiji (version 2.3.0) (Schindelin et al., 2012). After imaging, cells were digested with proteinase K (Thermo Fisher Scientific), and the nucleic acids, including the generated biotinylated cDNA, were recovered by phenol-chloroform extraction and concentrated by ethanol precipitation. RNA was digested with RNase H (NEB) and RNase A/T1 (Thermo Fisher Scientific) at 37 °C for 1 h, followed by biotinylated cDNA enrichment using Dynabeads MyOne Streptavidin C1 (Thermo Fisher Scientific). The 3′ cDNA adapter was ligated by T4 RNA ligase 1 (NEB) by incubating at 25 °C for 16h, and cDNA was recovered with the elution buffer of 95 % (v/v) formamide and 10 mM EDTA (pH 8.0) by boiling at 95 °C for 10 min, followed by ethanol precipitation. The library can be obtained by PCR amplification with NGS sequencing primer and gel purification of size between 180 bp and 400 bp. Sequencing was performed at the University of Chicago Genomics Facility on an Illumina NovaSeq 6000 platform in single-end mode with 100 bp.

### Sequencing data analysis

#### PolyA RNA-seq and nuclear RNA-seq

Raw RNA-seq reads were trimmed with Cutadapt (version 4.6) (Martin, 2011). The reads were first aligned to the human rRNA using STAR (version 2.7.10a) (Dobin et al., 2013) to further remove the rRNA contamination. The remaining unmapped reads were mapped to the human genome (GRCh38) with GENCODE v39 gene annotation using STAR (version 2.7.10a). Reads were assigned to gene regions using featureCounts (Liao et al., 2014). nTPM was calculated by RSEM (version 1.2.28) (Li and Dewey, 2011) and fold changes between different conditions were calculated by DESeq2 (version 1.38.3) (Love et al., 2014). Intron retention events were assessed using IRFinder (version 1.3.0) (Middleton et al., 2017) with default settings.

#### ARTR-seq

FastQC was used to assess the raw single-end FASTQ files. Cutadapt (version 4.3) was used for adaptor trimming. Reads were first mapped to the human rRNA using STAR (version 2.7.10a) to further remove rRNA contamination. The remaining unmapped reads were aligned to the human reference genome (GRCh38) using STAR (version 2.7.10a). Only alignments with at least 24 matched bases were included for downstream analysis. Mapped reads were deduplicated using UMI-tools (version 1.1.1) and counted with featureCounts (version 2.0.1) (Liao et al., 2014). For gene-level I_NSE_ calculation, total reads per gene were calculated by the sum of reads mapped to introns and exons for each RNA-seq library. Bioconductor package DEseq2 (version 1.38.3) was then used to perform differential analysis and calculate the speckle enrichment score (I_NSE_) (Love et al., 2014). In analysis Method 1, DEseq2 analysis was performed between N_SON_ (or N_SRRM2_) and N_-priAB_. In analysis Method 2, N_SON_ (or N_SRRM2_) was first subtracted by sequencing depth corrected N_-priAB_, as N_SON_corrected_ (or N_SRRM2_corrected_) = N_SON_ (or N_SRRM2_) –F_c_·N_−priAB_, where F_c_ is the correction factor calculated by ratio of total mapped reads of ARTR-seq using SON or SRRM2 antibody to the total mapped reads of ARTR-seq without primary antibody. DEseq2 analysis was performed between N_SON_corrected_ (or N_SRRM2_corrected_) and N_nu-RNA_. For intron-level and exon-level analysis (I_NSE(intron)_ and I_NSE(exon)_), mapped reads were first assigned to intron and exon regions based on “Ensembl_canonical” exons. Regions between two successive canonical exons were defined as canonical introns. DEseq2 was again used to calculate I_NSE(intron)_ and I_NSE(exon)_.

#### GO analysis

The functional enrichment analysis was performed using g:Profiler (version e110_eg57_p18_ 4b54a898) with g:SCS multiple testing correction method applying significance threshold of 0.05 and using Gene Ontology release 2023-07-27 (Kolberg et al., 2023).

#### Regression model

To identify the association of sequence features with enrichment, we used the regression coefficients of a generalized additive model (GAM), as computed using the pyGAM library (Servén and Brummitt, 2018). The regression is given by the equation:

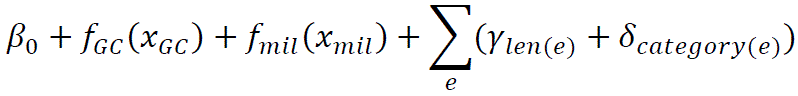

where β_0_ is a constant bias term, *f*_*GC*_ is a learned spline function applied to the gene’s GC content (*x*_*GC*_), *f*_*mil*_ is a learned spline function applied to the base-2 logarithm of the gene’s mean intron length (*x*_*mil*_), the sum runs over all internal exons *e*, γ_1_, …, γ_7_ are scalar coefficients used to score the binned exon length (*len*(*e*)), and δ_1_, …, δ_81_ are scalar coefficients used to score the exon category (*catego*r*y*(*e*)). The exon category is obtained by quantile binning and combining four values: MaxEnt 3’ splice site score, MaxEnt 5’ splice site score (Yeo and Burge, 2004), exon sequence ML score, and ±100nt flanking intronic sequence ML score (specifically, upstream from –120 to –21 and downstream from +6 to +105) (Liao et al., 2023). Since the ML model was trained on exons of fixed length, it is unable to account for differences in exon lengths properly; therefore, instead of using the raw score, we used the linear regression residual of the score with respect to exon length. Our intron-centric analysis was performed similarly, the only difference being the use of an intron category instead of the exon category. Intron category is computed similarly to exon category, the only exception being the use of the ML score of the ±100nt flanking exonic sequences instead of the exon sequence.

### Fluorescence labeling of FISH probe

DNA oligonucleotides were purchased from Integrated DNA Technologies (IDT). To prepare fluorescence-labeled probes, the 3’-end of oligonucleotides were first labeled with amine group as previously described (Gaspar et al., 2017). Briefly, to conjugate an amino-ddUTP at the 3’ end of each oligonucleotide, 66.7 μM DNA oligonucleotides, 200 μM Amino-11-ddUTP (Lumiprobe) and 0.4 U/μL Terminal Deoxynucleotidyl Transferase (NEB) were mixed in 1× Terminal Transferase Reaction buffer and incubated at 37°C overnight. The reaction was cleaned up by P-6 Micro Bio-Spin Column (Bio-Rad). For fluorophore labeling, amine-modified DNA oligonucleotides were mixed with 25 μg Alexa Fluor 647 (AF647, Thermo Fisher Scientific) or CF568 (Sigma Aldrich)-conjugated succinimidyl ester in 0.1 M sodium bicarbonate (pH 8.5) and incubated at 37°C overnight. The probes were cleaned up by ethanol precipitation and P-6 Micro Bio-Spin Columns. The labeling efficiency was calculated using NanoDrop One Spectrophotometer (Thermo Fisher Scientific #ND-ONE-W). The average probe labeling efficiency was ∼90%. The detailed sequences are provided in Table S2.

### Fluorescence labeling of antibodies

The secondary antibodies against mouse (Jackson ImmunoResearch, #715-055-150) and rabbit (Jackson ImmunoResearch, #711-005-152) were labeled with AF488 succinimidyl ester (Thermo Fisher Scientific). In brief, 24 μL antibodies (1 mg/mL) were mixed with 3 μL 10× PBS and 3 μL sodium bicarbonate (1 M, pH 8.5), 1 μL of DMSO dissolved AF-488 succinimidyl ester (1μg/μL) was added to the reaction and incubated at room temperature for 1 h. The labeled antibodies are purified by P-6 Micro Bio-Spin Columns (BioRad).

### RNA FISH

RNA FISH probes were designed using Stellaris probe designer and labeled as described above. After removing the medium and washing once with 1× PBS, the cells were fixed with 4% paraformaldehyde (PFA, Electron Microscopy Sciences) in 1× PBS at room temperature for 10 min. Cells were washed three times with DPBS and permeabilized with a solution containing 0.5% Triton-X100 (Thermo Fisher Scientific) and 2 mM vanadyl ribonucleoside complexes (Sigma-Aldrich, #R3380) in 1×PBS on ice. Cells were washed three times with DPBS, once with 2× SSC and once with wash buffer (10% formamide (Ambion, #AM9342) in 2× SSC). Cells were then incubated with FISH probes in hybridization buffer (10% formamide, 10% (w/v) dextran sulfate, 10 mM DTT in 2× SSC) at a final concentration of 1 nM per probe for 16 h at 37 °C in the dark. After hybridization, cells were washed twice with wash buffer for 15 min at 37 °C before being used for following immunostaining or imaging.

### DNA FISH

#### Cas9 targeting site and probe design

The Cas9 binding site against *LAMA5* genomic region was designed using CRISPR Guide RNA Design Tool using Benchling. To avoid interference with RNA FISH, the anti-sense strand was used for designing guide RNA and DNA FISH probes. The average spacing between each Cas9 binding site is 300 bp. Template DNAs with T7 promoter region for generating crRNAs were purchased from IDT. We designed DNA FISH probes by loading sequences between adjacent Cas9 binding sites into Oligoarray 2.1 (Rouillard et al., 2003) with the following conditions: length:18nt to 30nt; Melting temperature (Tm): 72 °C to 90 °C; GC content: 30% to 70%; Tm threshold for secondary structure formation: 54 °C; Minimal Tm to consider cross-hybridization: 54°C; prohibited sequences: GGGG; CCCC; TTTTT; AAAAA; the minimum distance between the 5′ ends of two adjacent oligonucleotides: 30; the maximum number of oligonucleotides to design per input sequence: 30; maximum distance between the 5′ end of the oligonucleotide and the 3′ end of the input sequence: 1000. Probes with multiple BLAST alignments were then removed to avoid non-specific binding. Designed probes were purchased from IDTDNA and labeled with AF647 using the same protocol as shown in Fluorescence labeling of FISH probe section.

#### Preparation of guide RNAs

crRNAs were synthesized using HiScribe T7 Quick High Yield RNA Synthesis Kit (NEB) according to the manufacturer’s instructions. All crRNAs are transcribed together, to make the transcription efficiency same for different crRNAs, we used a 10-nt common region to 5’-end of each crRNA to make the transfection efficiency homogeneous (Kocak et al., 2019). The synthesized crRNAs were purified using polyacrylamide gel electrophoresis. The guide RNA was assembled using 1:1 ratio of purified crRNAs and the Alt-R CRISPR-Cas9 tracrRNA from IDT in Nuclease-Free Duplex Buffer (IDT), incubated at 95°C for 5 min and slowly cooled down to room temperature over 1 h.

#### GOLD FISH and RNA FISH

To simultaneously detect DNA and RNA, we adapted the previously published GOLD FISH protocol (Wang et al., 2021). Briefly, cells were first fixed using prechilled MAA solution (methanol and acetic acid mixed in 1:1 ratio) at –20°C for 20 min and washed 3 times with 1× PBS and once with Blocking-binding (BBB) buffer (20 mM HEPES pH 7.5, 100 mM KCl, 5 mM MgCl_2_, 5% (v/v) glycerol, 0.1% (v/v) TWEEN-20, 1% (w/v) BSA with fresh added 1 mM DTT, 0.1 mg/mL *E.coli* tRNA and 1 U/μL RNaseOUT). After fixation, Cas9_H840_ (kind gift from the laboratory of Dr. Taekjip Ha) was assembled with annealed guide RNA in 1:1.4 ratio for 10 min at room temperature in BBB buffer. The cells were then incubated with assembled Cas9 RNP for 1 h at 37 °C. After incubation, 300 nM Rep-X (kind gift from the laboratory of Dr. Taekjip Ha) in BBB buffer supplemented with 2 mM ATP was added to cells and incubated at 37°C for 45 min. The cells were washed 3 times with 1× PBS, once with 2× SSC and once with 1× wash buffer. Cells were then incubated with DNA FISH probes (1 nM per probe) and RNA FISH probes (1 nM per probe) in hybridization buffer supplemented with 1 U/μL RNaseOUT for 4 h at 37 °C in the dark. After hybridization, cells were washed twice with wash buffer at 37 °C for 15 min.

### Immunofluorescence staining

After DNA and RNA FISH or RNA FISH alone, cells were fixed again with 4% PFA in DPBS for 10 min, washed 3 times with DPBS and blocked with 1mg/mL ultrapure BSA (50 mg/mL, Invitrogen) in DPBS for 30 min. Cells were immunostained with rabbit anti-SON antibody (1:200 dilution, Novus) or mouse anti-SRRM2 antibody (1:200 dilution, Sigma Aldrich) for 1h at room temperature followed by 3 times wash with DPBS. Cells were then incubated with AF488-labeled secondary antibody (1:200 dilution) for 1h at room temperature and washed 3 times again with DPBS.

### Fluorescence imaging

Before imaging, nuclei were stained with DAPI for 10 min and washed once with DPBS before imaging. To reduce photobleaching, 100 μL of imaging buffer containing Tris-HCl (50 mM, pH= 8), 10% glucose, catalase (67 μg/mL, Sigma Aldrich) and glucose oxidase (0.5 mg/mL, Sigma-Aldrich) in 2X SSC was used prior to imaging. For RNA FISH with immunostaining, imaging was performed on a Nikon Ti2-E inverted confocal microscope (Nikon AX-R) using either a CFI Plan Apo objective (60x oil, NA 1.40, Nikon) and GaAsP PMT detectors (DUX-ST detectors, Nikon). The pinhole size was maintained at 2 AU. Sample excitation was performed using the AS405/488/561/640 laser unit (LUA-S4, Nikon) with appropriate laser and filter settings. Z-stacks (0.2 µm step size, 7 stacks) were taken for each channel and an AI-based denoising (Nikon NIS-Elements AR 5.41.02 software) was applied. For GOLD FISH and RNA FISH with immunostaining, imaging was performed using a Nikon TiE microscope with a CFI HP TIRF objective (100x, NA 1.49, Nikon), and an EMCCD (Andor, iXon Ultra 888). Samples were excited with 647 nm laser (Cobolt MLD), 561 nm laser (Coherent Obis), 488 nm laser (Cobolt MLD) and 405 nm laser (CL2000, Crystal Laser).

### Image Analysis

Nikon NIS-Elements software (AR 5.41.02), Fiji ImageJ2 and MATLAB R2022b were used for image analysis.

#### Quantification of R_NS/NU_

Images were first denoised by NIS-Elements, Fiji was then used for maximum intensity projection and channel splitting. Individual cells were manually selected and saved as.tif files, which were subsequently analyzed in MATLAB with customized codes. The nuclei were segmented in the DAPI channel by Otsu’s algorithm, and nuclear speckles were segmented in the SRRM2 channel based on a global intensity threshold. Single-cell R_NS/NU_ values were then calculated by determining the mean RNA fluorescence intensity in nuclear speckles and dividing it by the mean intensity in the nucleus.

#### Quantification of GOLD FISH

Individual nuclei were manually selected in Fiji and saved as.tif files, followed by automated analysis in MATLAB with customized codes. For the DNA channel, a difference of Gaussians (DoG) filter was first used for background subtraction. Subsequently, DNA foci were identified by applying a global intensity threshold based on the mean and standard deviation of image intensity after the DoG transformation. A size threshold of 12 pixels was applied using MATLAB’s built-in function *bwareaopen* and the *regionprops* function was employed to extract centroid and area of each focus. RNA foci and nuclear speckles were identified using a similar approach, with a size threshold of 15 pixels. For each DNA or RNA focus, we calculated its center-to-center distance to the nearest nuclear speckle. This distance was then normalized by the sum of the radii of nuclear speckle and the RNA/DNA focus. An RNA or DNA focus was deemed NS-associated if the normalized center-to-center distance was less than 1.4. Similar association criteria were used for identifying class I RNA foci associated with DNA foci and class II RNA foci that were separated from the nearest DNA focus.

### RT-PCR

Total RNA was extracted using TRIzol™ Reagent (Invitrogen) according to the manufacturer’s instructions. DNA was removed using Turbo DNase (Invitrogen). RNA (1 μg) was reverse transcribed using iScript™ cDNA Synthesis Kit (Bio-Rad), and PCR was performed using Q5® High-Fidelity 2X Master Mix (NEB). The splicing efficiency is monitored using RT-PCR with primers located on two adjacent exons. The primer specificity is checked using Primer Blast. All primers are listed in Table S2. Amplified products were separated on a 1.5% agarose/TAE gel with Ethidium Bromide staining and visualized on a Bio-Rad ChemiDoc Imager. The unspliced efficiency is quantified by Fiji.

### Cell viability assay

Cell viability was assessed by measuring dead cells ratio stained with Trypan Blue using fluorescence microscope (Francis et al., 2020). Cells were seeded in an 8-well imaging chamber, after SON and SRRM2 double knockdown or treatment with control siRNA, the cells were stressed at 43 °C for 2 h. After heat shock, the cells were washed once with 1× PBS and stained sequentially with 1:10 dilution of Trypan Blue Solution (0.4%, Thermo Fisher Scientific) for 3 min and 1:500 of Hoechst 33342 (20 mM, Thermo Fisher Scientific) for 10 min. Imaging was performed on a Nikon Ti2-E inverted confocal microscope (Nikon AX-R) using a Plan Fluor objective (20x air, NA 0.50, Nikon) and GaAsP PMT detectors (DUX-ST detectors, Nikon). Cell viability was measured using Stardist as a plugin in Fiji (Schmidt et al., 2018).

