## Supplementary Table S1 for "Dynamics of RNA localization to nuclear speckles are connected to splicing efficiency"

**Table S1 - Related To Figure 3 - GOLD FISH analysis showing the co-localization of DNA, RNA and nuclear speckle**

| | % of NS-associated DNA | % of NS-associated RNA | % of NS-associated RNA with DNA associated in the same NS |
|---|---|---|---|
| rep1 | 0.722222222 | 0.689655172 | 0.5 |
| rep2 | 0.708333333 | 0.62962963 | 0.323529412 |
| rep3 | 0.782608696 | 0.681818182 | 0.366666667 |
| Mean | 0.737721417 | 0.667034328 | 0.396732026 |
| Stdev | 0.039488937 | 0.03262956 | 0.091996802 |

| % of NS-associated DNA-attached RNA foci | % of NS-associated DNA-detached RNA foci |
|---|---|
| 0.888888889 | 0.6 |
| 0.777777778 | 0.6 |
| 1 | 0.611111111 |
| 0.888888889 | 0.603703704 |
| 0.111111111 | 0.006415003 |
