## Supplementary Table S2 for "Dynamics of RNA localization to nuclear speckles are connected to splicing efficiency"

### Table S2 - List of oligos used in this study

| **RT-PCR primers** | Sequence (5' -> 3') |
|---|---|
| COL7A1_exon16_17_FP | ACTGCCACAGACATCACAGG |
| COL7A1_exon16_17_RP | GAGTCACATGGACCGTCCTC |
| COL7A1_exon22_23_FP | CTATCACCTGGACGGGCTG |
| COL7A1_exon22_23_RP | ATGCTTGGAACACGAGGTGA |
| THOC6_exon11_12_FP | CACCTCCGATCCTCCACACC |
| THOC6_exon11_12_RP | GCTCAGCTGCCACTGGTTG |
| THOC6_exon9_10_FP | CCAAGGAGGTCCAGACGAT |
| THOC6_exon9_10_RP | CCAGTCGGAATCAGTTGCC |
| MYO1C_exon21_22_FP2 | AAGACCCTGTTTGCCACAGA |
| MYO1C_exon21_22_RP2 | GGAATTTCTGCCGCCAGTG |
| MYO1C_exon13_14_FP | GCCCGTCCAGTATTTCAACA |
| MYO1C_exon13_14_RP | TGTGGATGGTGCTTGACAGT |
| RPS21_exon2_3_FP | CAGCCCAGCCTCGAAATGC |
| RPS21_exon2_3_RP | GGCACCGATGATGCGATTG |
| TUBB4B_exon3_4_FP | CCGTGCTCGTGGATCTGG |
| TUBB4B_exon3_4_RP | CCTTCTGTGTAGTGCCCCTT |
| KRT18_exon5_6_FP | GACGACGCTCACAGAGCTGA |
| KRT18_exon5_6_RP | CCCGTTGAGCTGCTCCATCT |
| NCL_exon13_14_FP | TGGACTGGGCCAAACCTAAG |
| NCL_exon13_14_RP | TCTTTCCTTGTGGCTTGTGGT |
| CALR_exon2_3_FP | GACTTCCCGCTGGATCGAAT |
| CALR_exon2_3_RP | GCCGACAGAGCATAAAAGCG |
| RFX5_exon7_8_FP | CAGCCAACTTTGGCAAGATCA |
| RFX5_exon7_8_RP | GGTGGCATAGACACCAAGGTC |
| TPI1_exon3_4_FP | GGCATGTCTTTGGGGAGTCA |
| TPI1_exon3_4_RP | CAGGCGATTACTCCGAGTCC |

| **Template DNA for *in vitro* transcription of crRNAs** | Sequence (5' -> 3') |
|---|---|
| *LAMA5* guide-RNA set (Figure 3D) | ACAGCATAGCTCTAAAACCAGAATG |
| | ACAGCATAGCTCTAAAACCTGTAAA |
| | ACAGCATAGCTCTAAAACAGGTGCC |
| | ACAGCATAGCTCTAAAACTTTTCAG |
| | ACAGCATAGCTCTAAAACTCTTGGC |
| | ACAGCATAGCTCTAAAACGGGCAAC |
| | ACAGCATAGCTCTAAAACCAAGAAC |
| | ACAGCATAGCTCTAAAACCGGGAAT |
| | ACAGCATAGCTCTAAAACCAAGGCT |
| | ACAGCATAGCTCTAAAACCATGGAC |

| **GOLD FISH probes** | Sequence (5' -> 3') |
|---|---|
| | TGGTTCTTTCAGCCAGCAGC |
| | CGCCTAACACACGCTGGGCC |
| | AGGCAGCTGGGAAACACGCA |

CCCTAGGGTGGCAGTGCAGG
CACTCAGAGTGGTGCAGCTGA
TGCAACCGAGCAGCCCAACTT
GGACCCGCCCTTGCCTTCTGT
GGGTGGAGCTCAGACAGG
TTCGGATGTGGCTGGTGGG
AGTCACCTGAGCCGGAGGA
ACCCACCCAGGCACTTGTT
GCTTCAGGTGCTGAGCTCC
CTTGTCCTGGGAAATTAAGTGCC
CGCAGCTCGTGACAGCAGGGAGT
TGACTGGGAATCTTTCATGGAAC
CACCCACCAGTGTCCCAAAACAT
CACCATCACCTTACTCCCACCCAA
ACTACCAGCCCACTCCCACCATCA
CCCTCTTGAATGATAGTTGAGCTG
GGATTCGTGGTTGATAGCTGCATC
CACTTTGGAGAAATGATTTGACTGT
CTCCAGGTGCTTGGGAGAGTCCAT
GCCATTGGTGTTCTGCAGATTTATT
CCATGTGTTCATTATTAATTCAAGA[illegible]
CATTCTTTCTGACTGTTGGCTCCCT
CCTTCTGGTTTCCTTCTCATAATTT[illegible]
CTCTCTCTTTATCTTCTGAGGTGTC
AGCTTCCTTTGCTCCTCTTCCACTT
AATCTCTCTTCCACTGTGTCTGTCT
TTAGCCCATCCATTGAGTCTTTAATT
GGCCTATGTTTTAGATTTCTTTATG
ACCTGCCACATTGTTTCTCTG
AGCAGGGTTTATCTTTGTGCTTCT
CCCTGGGATTGGGATGGTTTTGG[illegible]
CCAGCCCGGCACCACTTTTCCTAT
GAACACCAGTTCAGGCCCTTGCTT
CAGCTTCTTTGTTGATCACCTGTG
GTGGAGGACAGGACTAGAGC
CAGGGACTCCCGCTGTGGTCTGG
AGGTGCTTCCAACCTCAGGCCTT
TCCAGCTGTCACCCTGCCCACGG
GCAAGATTCCAGCCCTGTGACTT
AGCTGGAAGAAGGAAGCAGTTCT
CCTAGGGAGGTGACCAGGATGCC
TCTGGCTCAGGGAGGTTTT
CGAGTCCTGGCCGAAGTGG
TCAGTTTTGGAGAACGCAGGT
CCTGTGAGACCCTGTCCTCCA

AAGTTGCGTCAGAGCCACAGG
GCTGAGTCTGGGCGAGCGCTT
TTCAGGGAAGGGACTGAGC
ACAGGCTGCAGTGCACCTG
GCTTCAATCAGCAGCCGTG
GCCAACAGTGCCAACGAGT
TGCCTCCTCCCACACACGT
CCTGTGGAGGGCGTGCCAA

**T7 promoter template**

Sequence (5' -> 3')
GAAATTAATACGACTCACTATAGG

**RNA FISH probes**

Sequence (5' -> 3')
CATGAGGGACCTGGGTGC
TTCCCAGATGAGCTGAGG
ATGGGCACAGCAGCACTG
CTGTTTTCCACCACGGAG
CTGACAATGGGTGTGGGG
GGGTGACCAGCACAGAAG
AAGACAGCATGACACCCC
GACTCACAAAGCAGGTGG
TCACCCCTCAGGGATCAG
ACAGCTACTGAGCAGCTG
AACCAGTCTCCCATGAGG
GCAGGAGGAGGTACACAC
GGGGCACAAAGACCATGG
CAACCCTGTAGGGGATGG
GGAGCCTAGAACAGACCC
CAGGATGGCACCTCAGTG
CTACTTACGGGGTACGCC
CCAGCATCAGGCACACAG
CGACTTGTCCTGGCTGGG
CGGGGGAGAAAGGGTTGA
CGGGAGGACCGGATCATA
TGATCGCTGGGGCGGGAG
GAGGCCTGATGGTGAAGG
AAGGGTGTGATGGTCCGG
GGGTCTGATTGTAGTAGG
TGTGATCCTGGTATGGGG
TGATGGTGAAGGTGGGGG
TGTGATCGTGAGGTGGGG
CCATGAGGGGGGACTGAT
CATTGGTGGTGGTGTCTA
CTGCTCTGACGGCTGGTG
CGAGGGACCTGCAGACTC

CTGCTGTGTCTCCTCCAG
CTAAGCCCTCACTTCACT
AGAAGGGGTCCGTGACTG
CTGCAGTTGGAGGCATCC
GGAGAGGGGAGAGATGGC
CGAGTGGGCACAGAGACG
GCACAGAGACGAGGGGTG
ACAGACGGGTGGCGAGTG
GCACAGAGACGAGGGGTG
CACAGACGAGGGGTGGCG
GACGAGGGATGGCGAGTG
TGGCGAGTGGGCACGGAG
GTGGGCACAGAGACGAGG
GAGTGGGCACAGAGACGG
GCATGGAGAGACGGGTGG
GACGAGGGGTGGCGAGTG
TGGCGAGTGGGCACGGAG
GGGCACGGAGAGATGAGG
GAGAGATGAGGGGTGGCG
GATGAGGGGTGGCGAGTG
TGGCGAGTGGGCACGGAG
TGAGGGCAGAGGCGAGGG

| **RNA FISH probes** | Sequence (5' -> 3') |
|---|---|
| *LAMA5* | TCCGTATGCCGGAAGTTC |
| | TCGCGGGACACAGTGTTG |
| | CAGCACCATCATGAGCTC |
| | GATCTGCAGCTGCTCCAG |
| | CGCAGGAAGACAGCCGAG |
| | CACAGCTCCACATTGCTG |
| | CCTTTGACGTCCCGATAG |
| | ACATCGGCCCAGGAAGAG |
| | CCATGGCACTGACAAGGG |
| | GAGGCAGCGGTCTGAGTG |
| | TTCGGTGTTGTGCTGGCA |
| | CTGGCAGCGCTCACAGTG |
| | TGCTGCTCACGAAGCCAG |
| | CTGACACAGGGGGCGCTG |
| | AGGTTTGCAGAGGCACTG |
| | CAGGAGGCACCTGCATAA |
| | AAGAATCCGGGCGCACAC |
| | CATGGCTGGCAGGAGCTG |
| | AAGTTGGGGTCACCGTTG |
| | GTCGCAGTCGCTGAAGAG |
| | CACAGATCTCGCAGCGGG |

| | |
|---|---|
| | AGGGCGTTGCCGTAGAAG |
| | ACATGGGGTACAGTCGCA |
| | CGGTCCACAAGCACACGG |
| | TGGTCGGCAGTGGCACTG |
| *COL7A1* | CAGTGTCTGGCTGGACTC |
| | TGGCAGTAGAGTCTGGGG |
| | TGCAGCCCTGTGATGTCT |
| | CTGGTAGGTGGTTCCAGG |
| | CTCGCAGTACCGACACAG |
| | ACGATGACTGCAGCAGGG |
| | TGAGTCACATGGACCGTC |
| | GACAGATGAGCTGCTGGC |
| | AACCCTGGTCCAGGTAAT |
| | TGTATCCTGTGGCGCCAG |
| | GGCTGAGTGCCAGGAAAC |
| | CAGTCCATCCAGCTCAGC |
| | GTATACTCAGTATCTGGC |
| | CACATGGGCCCTCACATG |
| | ACAACCACAGAGGCAGGG |
| | CACGACCCACAGGCTCAG |
| | CGTCGCTGGAAGCATTGA |
| | TACCCAGGTGATCCGTAG |
| | GTCTGTAAGCTGTGGCTC |
| | AGTATCTGGTGCCTCATG |
| | AGAGTCTGTGTTTCCTGG |
| | TTCGAGACCCCGGATCTC |
| | CAAGTGCAGTCACTCGCA |
| | AATGGAGACAGGTGTGCC |
| | TGCACCACGTGAAGCGTC |
| *HMG20B* | CGCGACGCTCCAACCAAC |
| | AGAAACTTTCCCGGACCT |
| | TTGGGGCCGTGGGACATG |
| | CGCTCTTGCTTGACAGTC |
| | CTTCTTCCGCTTCTTGCC |
| | TGGGCCCATTCGGCAGAA |
| | TCGTTCAGGAAGCGCACG |
| | AAAGGGCAGATCCGGGTG |
| | CCAGCATCTTGGTGATCT |
| | TGCAGCTTGCTCCACTCG |
| | CCGCAGCTCCTTCATGTA |
| | CTTCAGACTGCTGGTACG |
| | TCCGTGCACATCTTATAG |
| | CTTCTTCTCCTGGATCTT |

| | |
|---|---|
| | GTGTCCATTCAGGAGAGT |
| | TGGAGAAGCCATCGCAGT |
| | ATTCATCTTCCGCAAGCG |
| | CGTTCTGCTCCTCGAAGG |
| | CTGCTCATGCTCTGCGTG |
| | GCAGTGAGGCGAAGCTGG |
| | GGCCATGTAGAAGTCCAG |
| | CTCGATGGCTCCGTGAAG |
| | ATGAGCTTCTCGTGCTGG |
| | CCAGGATTTCCTTGATGC |
| | CACTCCTCACAGGTGCTC |
| *U2AF2* | GCACGTCCCAGTATTTAC |
| | ATGTGCTCAAAGCCTGGG |
| | CTTGTACTGCATTGGGGT |
| | AGAAGAGCAGTGGCTGGA |
| | GTCAGGGGTCATGGTGGG |
| | TTGGGGTCACAGCCAGAC |
| | CTTGTCTGGTCATCTGGC |
| | GATGTTGCCCACGTAGAG |
| | CGCATCTGGGCGTTGAAG |
| | CAACACTGGGTTGCCAGG |
| | GTCCTGGTTAATCTGCAC |
| | GGGTAGTCTCGTCCACTG |
| | CCATCAAAGGCCATAGCC |
| | CTGGCCCTGGAAGATGAT |
| | GCCTGCGGATCTTTAGTG |
| | AAGCGGCTGGTAGTCGTG |
| | GGTTCTCTGACATGCCAG |
| | GGGACCACAGTGGACACA |
| | GAACAGCTTGTGGGCAGA |
| | CAGGTAGTTGGGTAAGCC |
| | GCCCAAAGGATGTCAGCA |
| | AGGTTGAAGGCCTTGAGG |
| | CGTGGCACTGTCCTTGAC |
| | CTCACAGAAGGCGTAGCC |
| | TCCGTGACGTTGATGTCC |
| *COL27A1* | GCTGGAGGATGTCCACAT |
| | AAGGAATGACTCCCGGGG |
| | CCCGCTGCGTAAAGATGA |
| | AGGAAGGCATGGTTCACC |
| | CTGCAGCTTGCGTTTCTG |
| | AGGTGGACGACCGTCTTG |
| | AGGTCGAAGGCCACTGAG |

| | |
|---|---|
| | CACCAGAGTGACTGTGCG |
| | CCTGTGGAAAGGCAGCAG |
| | CAAAGAGGAAGGAGCCCC |
| | AACTGGACTGCATGCGGG |
| | AACTGGCAGAGAGCACCT |
| | CTGCGTCACAGGGTAGAT |
| | TTCCTCAGGTGGGTACAG |
| | AAGGTCTGCCAGAGTCTT |
| | GCGAGGTCGGACTGGAAG |
| | GTGGTCAAGTTCTCCAGG |
| | CTAGTCCTTTGGGGCTTG |
| | TGCTGGTGAGGGTTTGTG |
| | CAGCTTGGCAGGTAGCAG |
| | TCAAGTGCGTTACTGGCT |
| | ACAGAGGCTGGGAGCATG |
| | ACAGAGGCTGGGAGCATG |
| | CTGTGATGGTTGAGCGGC |
| | GGGATTTTGGTGGCTGTG |
| *ABCC2* | CCGTTGTCTAGGACCATT |
| | GCCGCACTCTATAATCTT |
| | GTAGCAGTTCTTCAGGGC |
| | GTAAAAGGGTCCAGGGAT |
| | CAGCTTCCTTAGCCATAA |
| | GTGCTGTTCACATTCTCA |
| | CTATCTCCAACCTGTCAC |
| | GCCATGCAGAATGCAGGA |
| | GTAAAGAGGCCTAGGTCC |
| | GGGTTATACCTGCAATGT |
| | CTGATGAACCTGTGTCAT |
| | CAGTGCAGACTAAAACCC |
| | ATAAAGCCAAGGTCTGCC |
| | CTGCCCAGGAGATCTTAA |
| | ACTTAGGCTGGCTTCTGA |
| | GAGGTGAGGAAGGCCCAA |
| | AACCTGACACAAAGGCCC |
| | CAGACCGTGAAATGCTGA |
| | ATGTTAAGCACCTCACTA |
| | GCCCTTCATTTCAGAAGT |
| | GGCTATGTGTAGGAGGCA |
| | CCATCCATGCTGTGATTC |
| | CCCAGCCAACACTTCAAT |
| | TTCTCCATGAATAGCGCC |
| | CTAGATCTGGTTTGCAGA |

| | |
|---|---|
| *NCL* | CAGCGAGAGCTCGAGACT<br>GAGCACGTACACCCGAAG<br>CCGCGGGTGCTGAAGATC<br>AAGCCAAGCGACGGCGAT<br>GCGCAGATGAGTCCAGAA<br>CGGAGTGTGAAGCGGACA<br>AGCTTCACCATGATGGCG<br>ATTTTTACCTGCCTTCGC<br>ATTTTCTTGGGGTCACCT<br>TTCTGAGGTATGACGACC<br>AGCAGCCTTCTTGCCTTT<br>CCTTCTTTGCTGAGGTTG<br>TTTGTTGGGGAAACGACC<br>TGTGGCAACTGCAACCTT<br>GACAGCTGCTTTCTTGGC<br>CTGCCTTTTTGCCTGGAG<br>TCTTCTTGGCAGGTGTTG<br>GCTTTGGCTGGTGTAACT<br>TACCAATGCTTTGCCTGG<br>CCTTCTTACCAGGAGTTG<br>CCGTCAAATGACTCCTTT<br>TATCCTTGCCCGAACGGA<br>CAGTTTCCCGGTCAGTAA<br>CCAAACCCTTTGGAGGAC<br>TCCTCACTGTTGAAGTCT |
| *SFPQ* | GCGGCGGAGAACGGAAGT<br>GCGATTCTGATTGAGGCC<br>GATCGGAGGCTTAGGGCC<br>GCAACGACGGGCTTGGAA<br>AGCTGGAGGCTGGTGGTG<br>GAGGTTGGTGGAGTGGCG<br>CGAGGTGACTGCAGGCGG<br>AGGAGGTGTGGTAGGGAC<br>ATTTTGCCGCCTTTGGGA<br>GAGATCTTCTCCTCGCTG<br>TTTAAACCCCTCCGAGTC<br>GCCTCCTCAAGAGAGACA<br>GTGTAAGTTTTCTCTCCA<br>CAAACAACCGACATCGCT<br>ATCAGCAGGTAGATTCCC<br>TGAATTCATCCTCCGTGA<br>CTCCTGGTTCTCCATATT<br>AATCCGAATCCTTTGCCT |

| | |
|---|---|
| | GCCAAAGCTCTAGATTCA |
| | CAGTTCGGCTTTGGCAAT |
| | GGTCTTTTTACCTTGGCG |
| | TTCTTCCTGTCGTCTCAT |
| | GTTCTTCCATGCGTCTTA |
| | CTCCTCTTGCCTCAATTG |
| | TCCATCTCACGTTGACGA |
| *P4HB* | CAGCGGGGGCGACGAGAG |
| | GCATGTCGGACACGGATC |
| | CGCGAAGTTGCTTTTCCG |
| | CACCAGCAGGTACTTGTG |
| | GCTTTGGCATACTCAGGG |
| | GAACCTTCTGCCTTCAGC |
| | CTTGGCCAACCTGATCTC |
| | AGGTCAGACTCCTCCGTG |
| | CTTGATGGTGGGATAGCC |
| | CGTGTCTCCATTCCTGAA |
| | GCTGTATATTCCTTGGGG |
| | ACTCCACCAAGGACTCTG |
| | AGAAGCCGATGACAGCCA |
| | CACACCAAGGGGCATAGA |
| | ATGTCATCAGCCTCTCTG |
| | CTTCTTCAGCCAGTTCAC |
| | GTCGAGCTGGTATTTGGA |
| | TCTGCTGCCTGCAAAAAC |
| | TGGTATGTCATCGATGGC |
| | AGAGGACAACCCCATCTT |
| | GTCTGGCTTTGCGTATTAC |
| | AGAGAGGTTCCCTGGGTTT |
| | TAGGGGTGAGGTGTCACTT |
| | CGGGGGTGAACGGACGGTG |
| | GGATCCCTTTCCAAAAACC |

AGCCCGGCCCTACTATTCATCCCTATAGTGAGTCGTATTAATTTC
CTGGAGCGTGGACTATTCATCCCTATAGTGAGTCGTATTAATTTC
GGTGAGTCAGGGGATATTCATCCCTATAGTGAGTCGTATTAATTTC
ACAATTCCGGATATATTCATCCCTATAGTGAGTCGTATTAATTTC
CTAACATGGTGAAATATTCATCCCTATAGTGAGTCGTATTAATTTC
GAGTGTGATACTCTTATTCATCCCTATAGTGAGTCGTATTAATTTC
CTCACAAAATGGATATTCATCCCTATAGTGAGTCGTATTAATTTC
TCAATACTGCTTTCTATTCATCCCTATAGTGAGTCGTATTAATTTC
TCAGATTTCGCTTATATTCATCCCTATAGTGAGTCGTATTAATTTC
CACGGCCCCCCGCTTATTCATCCCTATAGTGAGTCGTATTAATTTC

CT
TGT
GT
CATC
TTCCT
TATT
TTCTG
CACTA
GCTGT
TTTAT
ACCA

G
