## Supplementary Information for "Dynamics of RNA localization to nuclear speckles are connected to splicing efficiency"

**Supplementary Figures S1-S8**

**Supplementary Table S1:** GOLD FISH analysis

**Supplementary Table S2:** List of oligos used in this study

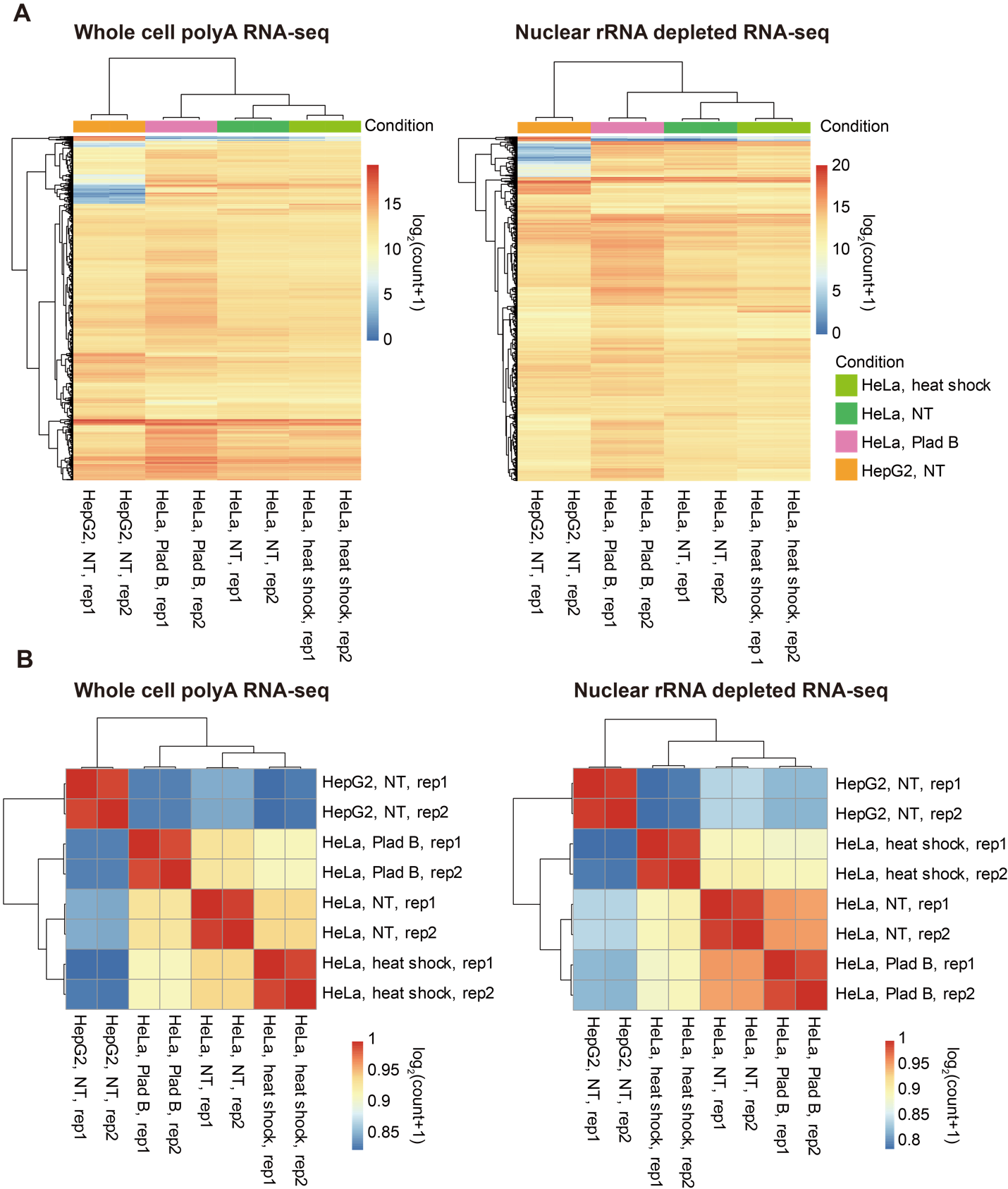

**Figure S1: Library quality of polyA RNA-seq and nuclear RNA-seq datasets.** (A) Heatmaps showing the expression of top 1000 genes (count sum) in polyA RNA-seq (left panel) and nuclear RNA-seq datasets (right panel), organized by hierarchical clustering based on log2-transformed counts (log_2_(count+1)). Gene counts were transformed using the normTransform() function from the DESeq2 software (v1.34.0) (Love et al., 2014). Figures were generated using the pheatmap package (v1.0.12) in R software (Kolde, 2019; R core team, 2022). Each experimental condition has two biological replicates. Scale bar: log_2_(count+1). (B) Heatmap showing the correlation between two samples from polyA RNA-seq datasets (left panel) and nuclear RNA-seq datasets (right panel) based on regularized log (rlog) transformed counts for genes calculated by the DESeq2 software (v1.34.0) (Love et al., 2014). Samples were organized by hierarchical clustering, and figures were generated using the pheatmap package (v1.0.12) in R software. Scale bar: Pearson’s correlation coefficient.

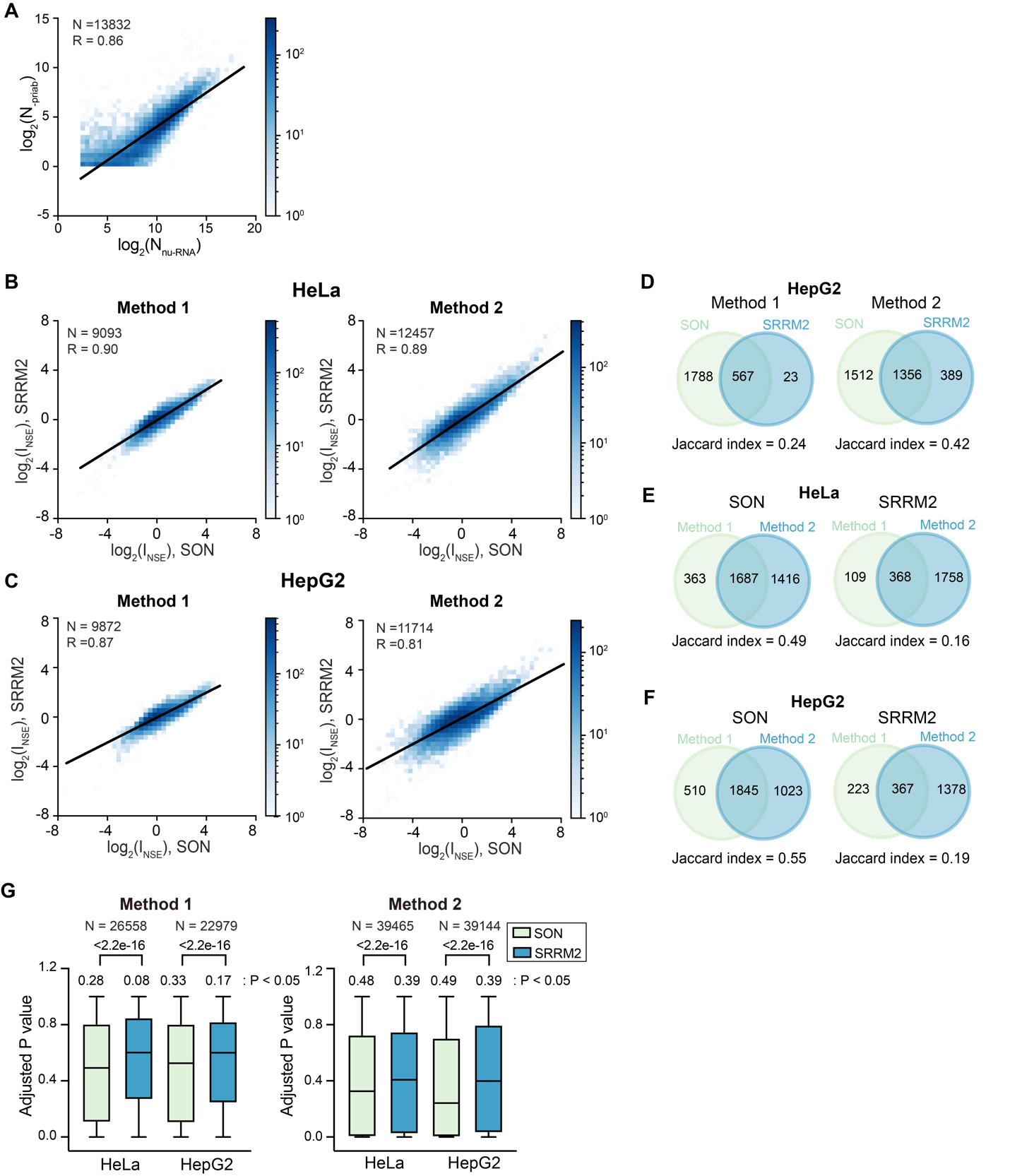

**Figure S2. Comparison between the choice of targeting proteins and analysis method.** (A) 2D histogram showing the correlation between mapped reads to each gene from ARTR-seq dataset obtained from sample prepared without primary antibody (N_-priAB_) and rRNA-depleted nuclear-RNA seq (N_nu-RNA_). (B)-(C) 2D histogram showing the correlation between I_NSE_ determined through targeting SON and SRRM2 in (B) HeLa cells using Method 1 and Method 2, and (C) HepG2 cells using Method 1 and Method 2. Genes with lfcSE<1 from DESeq analysis of ARTR-seq are included. “N” reports the total number of genes included in each plot, and “R” reports the Pearson’s correlation coefficient in (A)-(C). (D) Venn diagram of overlapped speckle-enriched genes identified through targeting SON and SRRM2 in HepG2 cells using Method 1 or Method 2. (E)-(F) Venn diagram showing the overlapping genes between Method 1 and Method 2 when targeting SON or targeting SRRM2 in (E) HeLa cells and (F) HepG2 cells. (G) Comparison of the adjusted P value for each gene in DEseq analysis of speckle enrichment between targeting SON and SRRM2 in HeLa cells. Both Method 1 and Method 2 shows less variations with SON antibody than with SRRM2 antibody. P-values calculated using unpaired t-test and number of genes included in each comparison (N) are labeled above each box plot. The fraction of genes with adjusted P-value less than 0.05 in each case is labeled above each box plot.

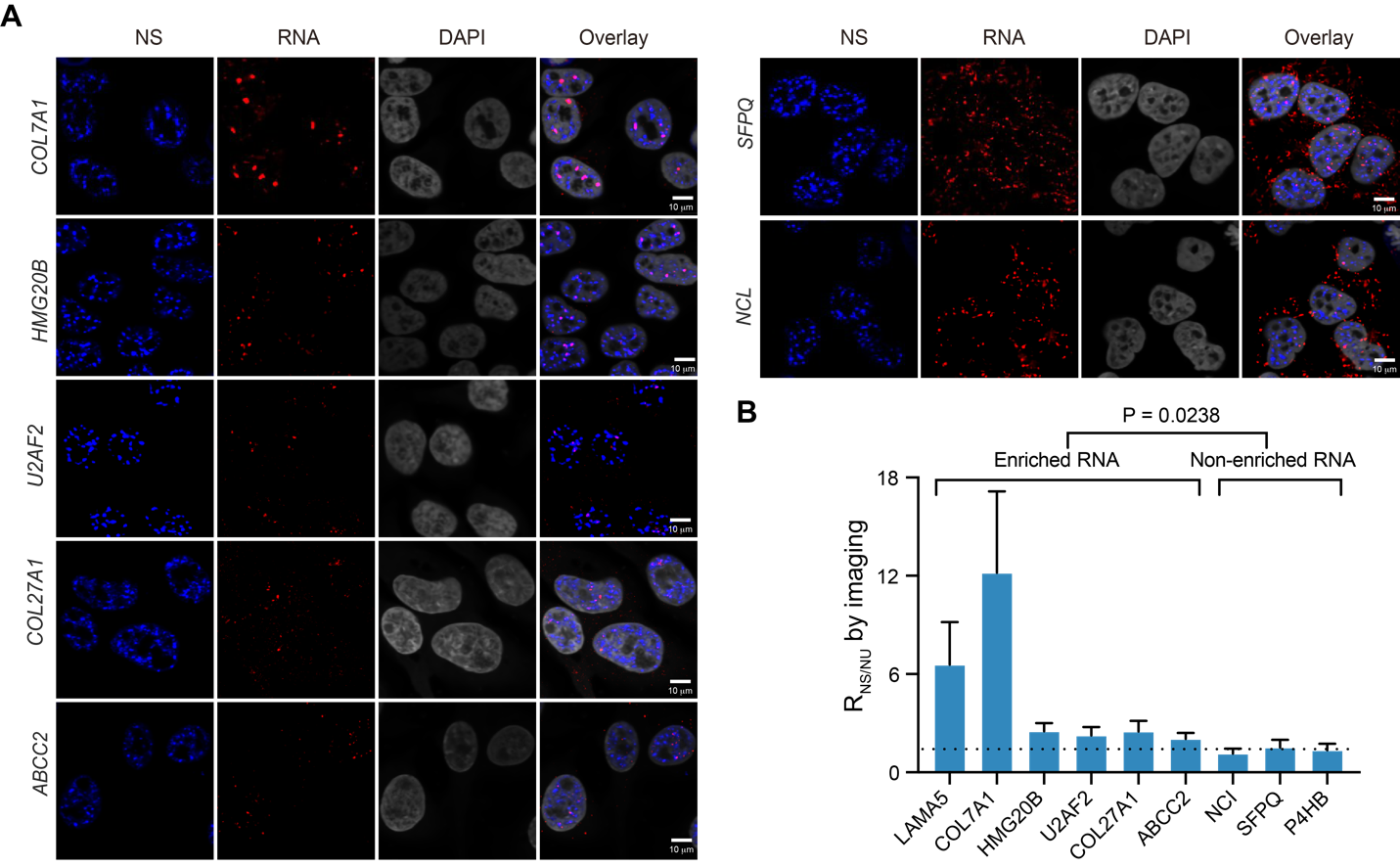

**Figure S3.** **Additional RNA FISH imaging validation of ARTR-seq on selected genes.** (A) RNA FISH images showing speckle-enriched *COL7A1*/*HMG20B*/*U2AF2* transcripts in Hela cells, speckle-enriched *COL27A1*/*ABCC2* transcripts in HepG2 cells and speckle non-enriched *SFPQ*/*NCL* transcripts in Hela cells. Nuclear speckles (NS) were stained with AF488 labeled antibodies against SON (blue), RNAs were labeled with AF647 labeled FISH probes (red), and nuclei were stained with DAPI (grey). Scale bar: 10 μm. (B) Comparison of R_NS_/_NU_ values determined by RNA FISH imaging in (A), between speckle-enriched transcripts and non-speckle-enriched transcripts classified consistently using Method 1 and Method 2 of ARTR-seq analysis. P-values were calculated using unpaired t-test.

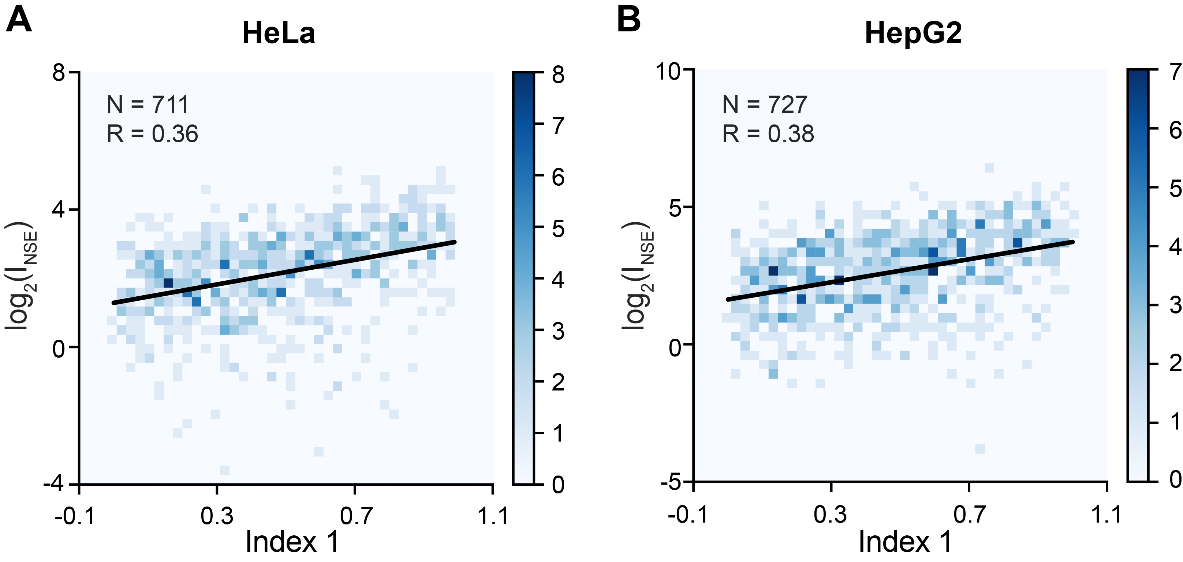

**Figure S4: Comparison between ARTR-seq and APEX-seq methods for nuclear speckle transcriptome analysis.** 2D histogram showing the correlation between I_NSE_ determined by ARTR-seq using Method 1 in HeLa cells (A) or HepG2 cells (B) and Index 1 from APEX-seq (Barutcu et al., 2022). A higher Index 1 indicates higher speckle enrichment. Genes with lfcSE<1 from DESeq analysis of ARTR-seq, and genes with Index 2 > 0.5 in APEX-seq are included. Index 2 in APEX-seq was used to evaluate the expression level change of genes upon expressing APEX-fused proteins. Total number of genes included in each plot is indicated by “N”, and Pearson’s correlation coefficient is reported by “R” value.

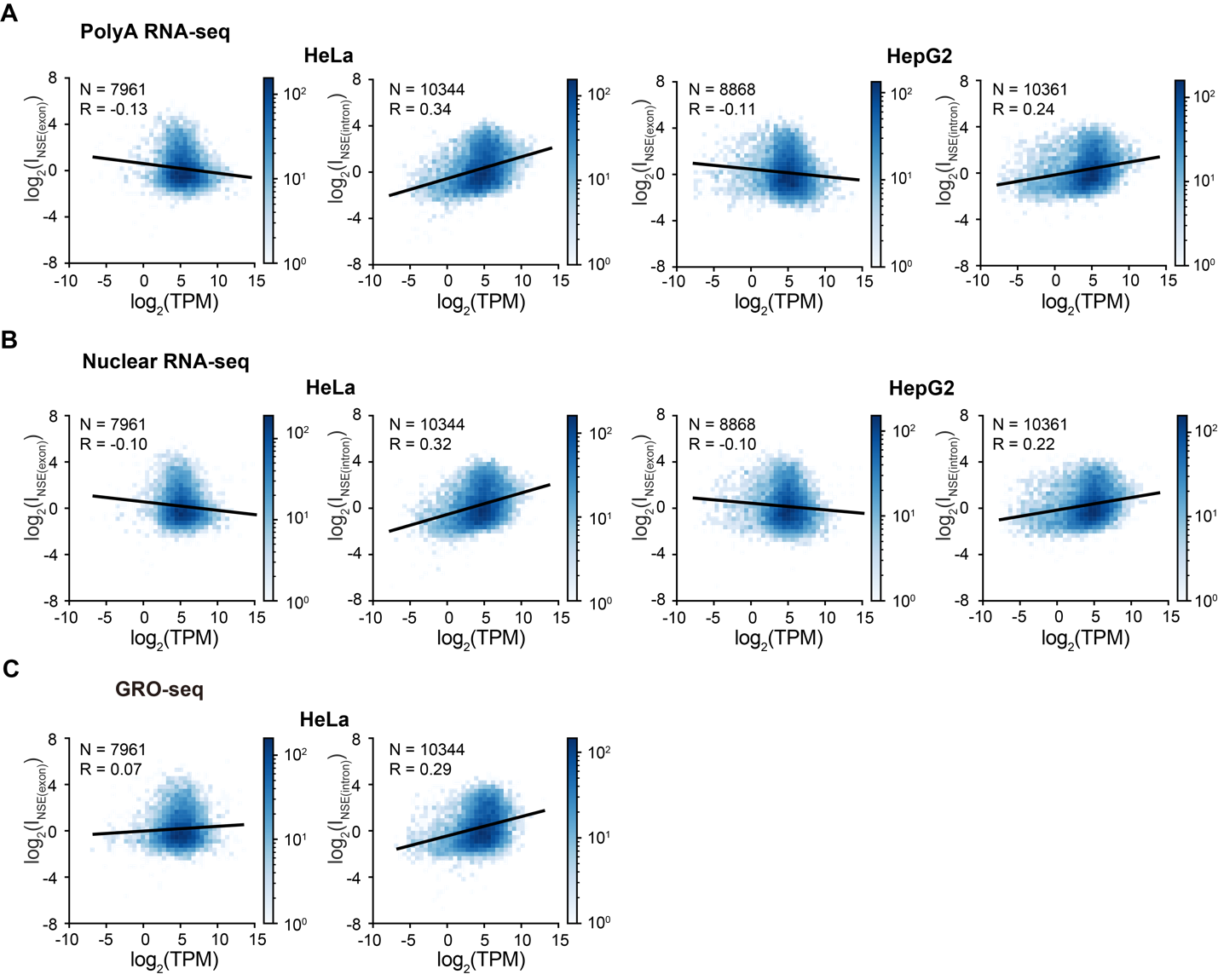

**Figure S5: Correlation between transcript speckle enrichment and transcript abundance.** 2D histogram showing the correlation between I_NSE(exon)_ or I_NSE(intron)_ determined by ARTR-seq using Method 1 and TPM (in log_2_ scale) from (A) polyA RNA-seq in HeLa cells or HepG2 cells; (B) rRNA-depleted nuclear RNA-seq in Hela cells or HepG2 cells; (C) GRO-seq in HeLa cells (Andersson et al., 2014). TPM: transcript per million reads. TPM was calculated by RSEM (Li and Dewey, 2011). Genes with lfcSE<1 from DESeq analysis of ARTR-seq are included. Total number of genes included in each plot is indicated by “N”, and Pearson’s correlation coefficient is reported by “R” value.

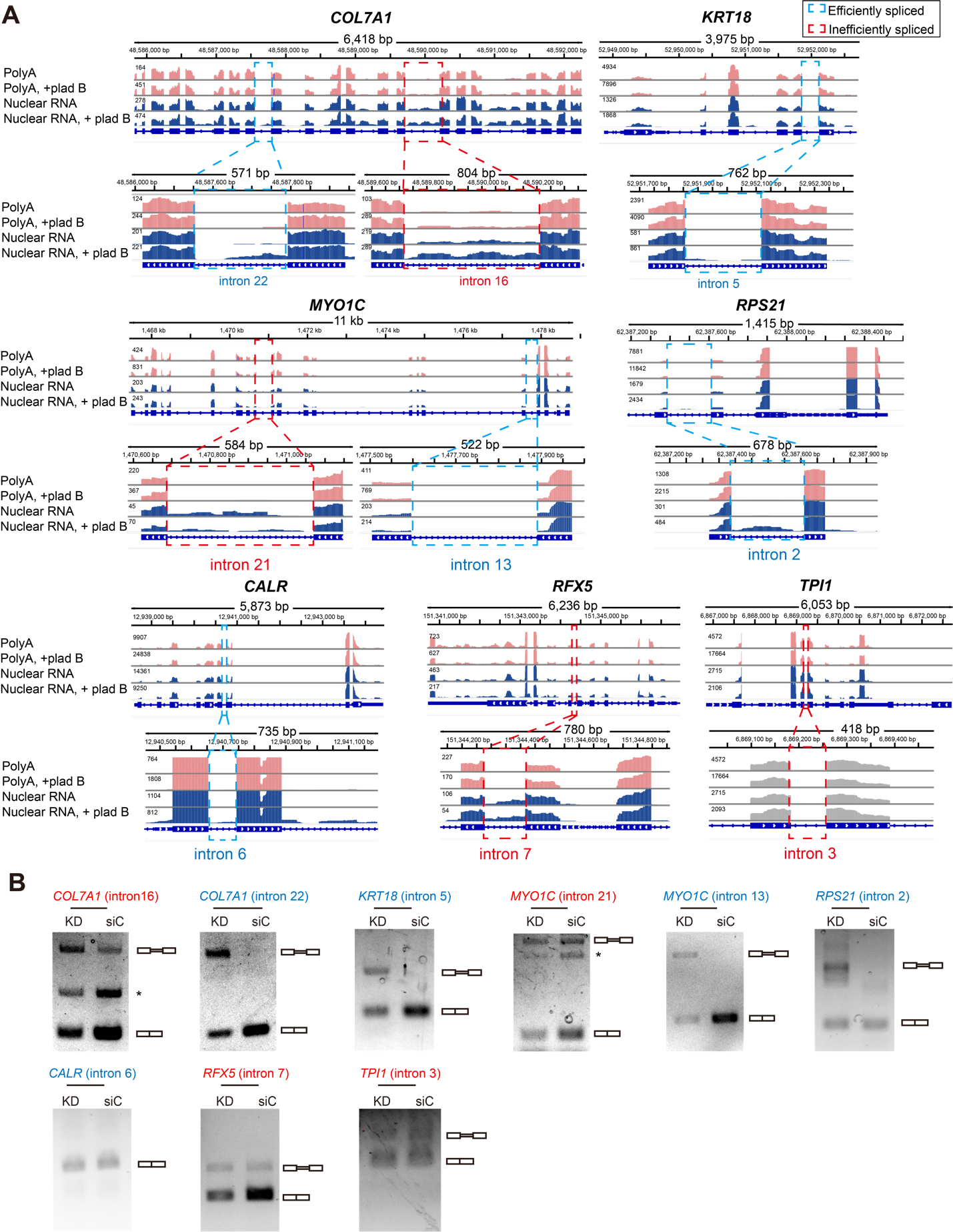

**Figure S6: Genome tracks and electrophoresis analysis of additional selected genes in RT-PCR assay.** (A) Genome tracks showing polyA RNA-seq (pink) and nuclear RNA-seq (blue) under NT and Plad B treatment conditions for additional selected Group A genes (*COL7A1*, *MYO1C*), Group B genes (*KRT18*, *RPS21*) and Group C genes (*CALR, RFX5, TPI1*). Selected introns are highlighted in cyan and red for efficiently spliced introns and inefficiently spliced introns respectively. (B) Electrophoresis analysis of RT-PCR products from Group A gene (*COL7A1*, *MYO1C*), Group B gene (*KRT18*, *RPS21*), and Group C genes (*CALR, RFX5, TPI1*), either upon double knockdown of *SON* and *SRRM2* (KD), or treatment of control siRNA (siC). Asterisks denote unknown bands, which were not considered in the calculation of apparent fraction of unspliced intron.

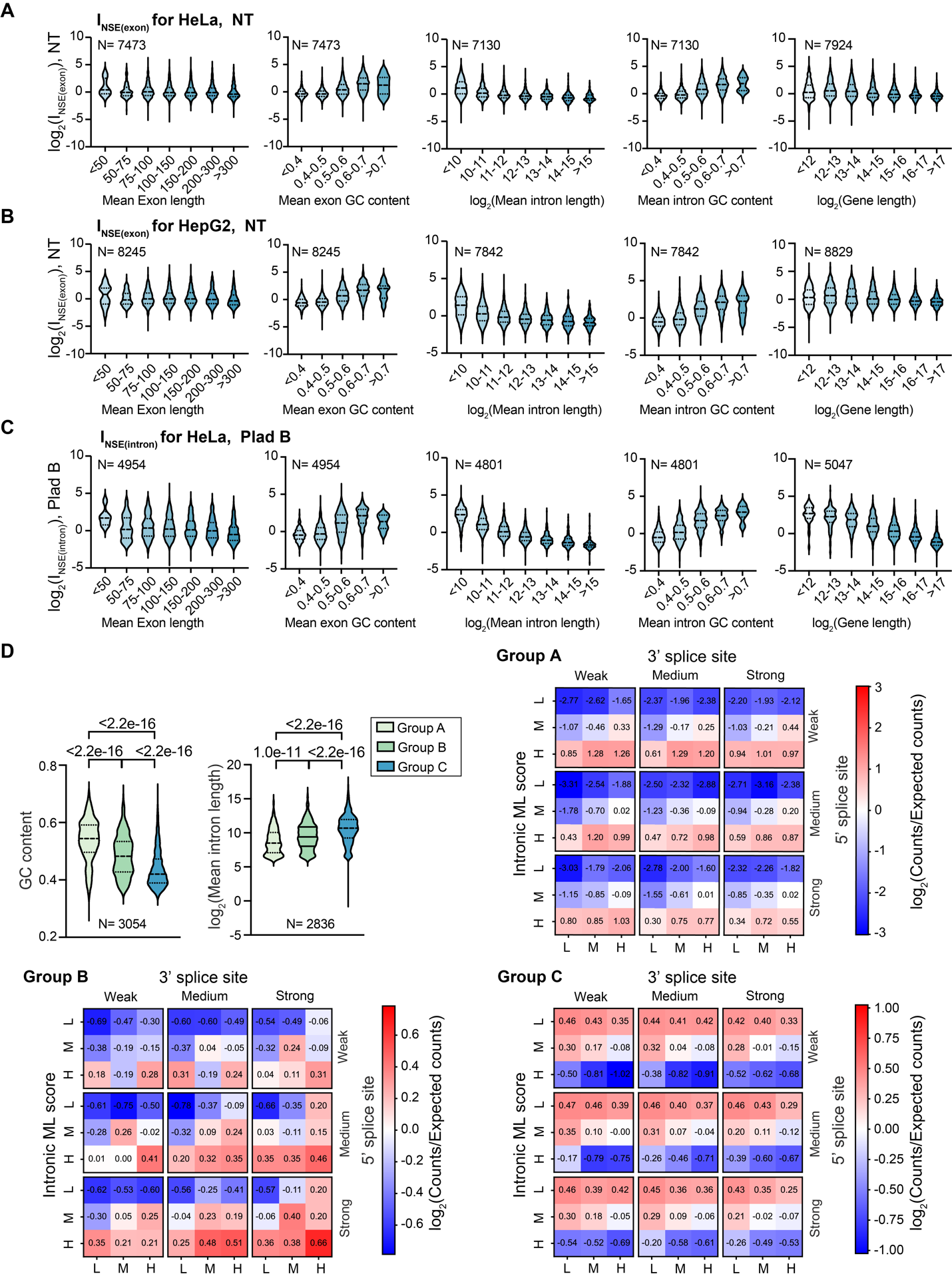

**Figure S7: RNA sequence features associated with speckle enrichment.** Comparison of transcript speckle enrichment in each bin based on mean exon length, mean exonic GC content, mean intron length (log_2_ scale), mean intronic GC content and gene length (log_2_ scale) of each transcript for (A) I_NSE(exon)_ HeLa cells under NT condition; (B) I_NSE(exon)_ in HepG2 cells under NT condition; (C) I_NSE(intron)_ in HeLa cells with Plad B treatment. Genes with lfcSE<1 from DESeq analysis of ARTR-seq are included. Total number of genes included in each plot is indicated by “N”. (D) Comparison analysis of GC content, intron length, and splicing-related features in Group A, B, and C transcripts. GC content and intron length of Group A, B and C genes are shown in violin plots. P-values were calculated using unpaired t-test. Splicing-related features represented as a combination of ML score with splice site strength for Group A, B and C genes are shown in heatmaps. The color bar indicates the log2 fold change of counts of each splicing-related combination compared to expected counts. Blue: a combination with lower-than-expected count, or depleted; red: a combination with high-than-expected count, or enriched.

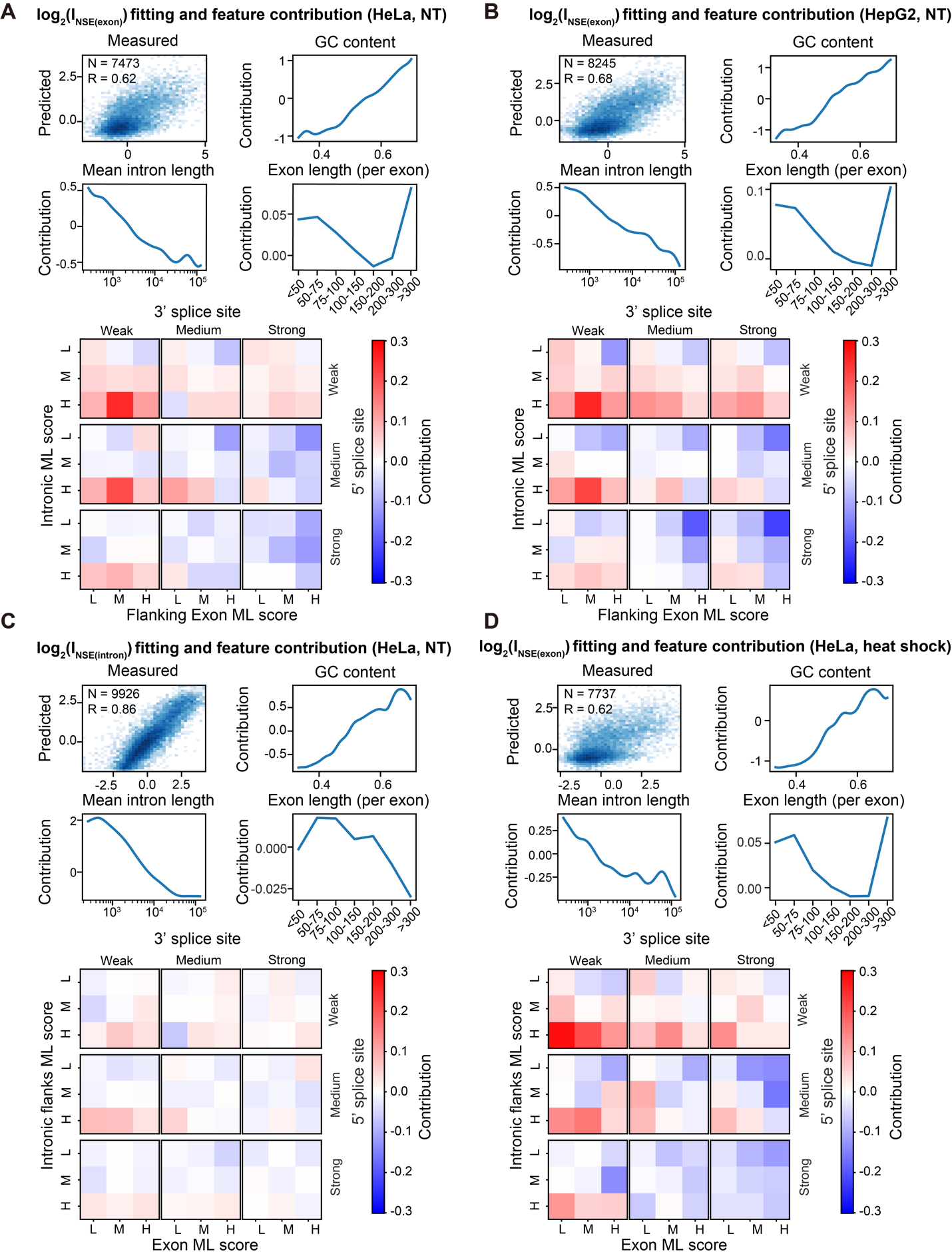

**Figure S8. Additional regression analysis on transcript speckle enrichment in different conditions.** Input parameters and other related details of the regression model are described in the main text and Methods. (A-B) The intron-centric regression model reveals contributions from GC content, mean intron length, individual exon length, and a combination of splice site strength, intronic ML score and flanking exonic ML score to I_NSE(exon)_ values in HeLa cells under NT (A), and in HepG2 cells under NT (B). (C) The exon-centric regression analysis on I_NSE(intron)_ values in HeLa cells under NT. (D) The exon-centric regression analysis on I_NSE(exon)_ values in HeLa cells upon heat shock.
